# TRPM2 ion channels steer neutrophils towards a source of hydrogen peroxide

**DOI:** 10.1101/2021.01.06.425587

**Authors:** Hassan Morad, Suaib Luqman, Chun-Hsiang Tan, Victoria Swann, Peter A. McNaughton

## Abstract

Neutrophils must navigate accurately towards pathogens in order to destroy invaders and thus defend our bodies against infection. Here we show that hydrogen peroxide, a potent neutrophil chemoattractant, guides chemotaxis by activating calcium-permeable TRPM2 ion channels and thus generating an intracellular leading-edge calcium “pulse”. The thermal sensitivity of TRPM2 activation means that chemotaxis towards hydrogen peroxide is strongly promoted by small temperature elevations, suggesting that an important function of fever may be to enhance neutrophil chemotaxis by facilitating calcium influx through TRPM2. Chemotaxis towards conventional chemoattractants such as LPS, CXCL2 and C5a does not depend on TRPM2 but is driven in a similar way by leading-edge calcium pulses. Other proposed initiators of neutrophil movement, such as PI3K, Rac and *lyn*, influence chemotaxis by modulating the amplitude of calcium pulses. We propose that intracellular leading-edge calcium pulses are universal drivers of the motile machinery involved in neutrophil chemotaxis.

## Introduction

The first line of defence against pathogen invasion is formed by tissue-resident “sentinel” macrophages that detect characteristic pathogen-associated molecular patterns, engulf the invading pathogens and then kill them, partly by generating a “respiratory burst” of toxic free radicals. Following interaction with pathogens, macrophages release a range of chemoattractant signals, including cytokines and chemokines such as IL1, TNFα and CXCL2. In addition, destruction of bacterial proteins releases chemoattractant signals, such as fMLP and activation of the complement cascade releases signals, such as C5a. In response to the release of these chemoattractants, circulating neutrophils leave the blood space and navigate up the chemoattractant gradients to locate and attack invading pathogens. A subsequent wave of macrophages and lymphocytes, such as natural killer (NK) T cells, joins the attack in a similar way^1^.

While these key features of the early response of the innate immune system have been known for some time, important questions remain. Are other chemoattractants also important? Hydrogen peroxide (H_2_O_2_) is a neutrophil chemoattractant *in vitro*^2^, and more recent *in vivo* evidence shows that H_2_O_2_, generated by the reaction with water of reactive oxygen species (ROS) produced by the NADPH oxygenase DUOX, forms a concentration gradient extending more than 100μm from a site of injury, and moreover that H_2_O_2_ is a critical early guidance signal for neutrophil chemotaxis *in vivo*^3–11^. How does H_2_O_2_ act as a neutrophil chemoattractant? More generally, what are the cellular signalling mechanisms that guide neutrophils up gradients of chemoattractant molecules? The present study aimed to elucidate the mechanisms that couple gradients of H_2_O_2_ and other chemoattractants to neutrophil chemotaxis.

## Results

### Hydrogen peroxide is a potent neutrophil chemoattractant both *in vivo* and *in vitro*

We used a range of methods, both *in vivo* and *in vitro*, to confirm that H_2_O_2_ is a neutrophil chemoattractant. H_2_O_2_ injected into the peritoneal space *in vivo* recruits mouse neutrophils, with efficacy observed down to a concentration of 10nM (Fig. 1A). Recruitment remains significantly elevated at concentrations between 10nM and 10μM but is inhibited at 100μM. The well-established neutrophil chemoattractant fMLP also potently recruits neutrophils, with a similar efficacy to optimal concentrations of H_2_O_2_.

**Fig. 1.**
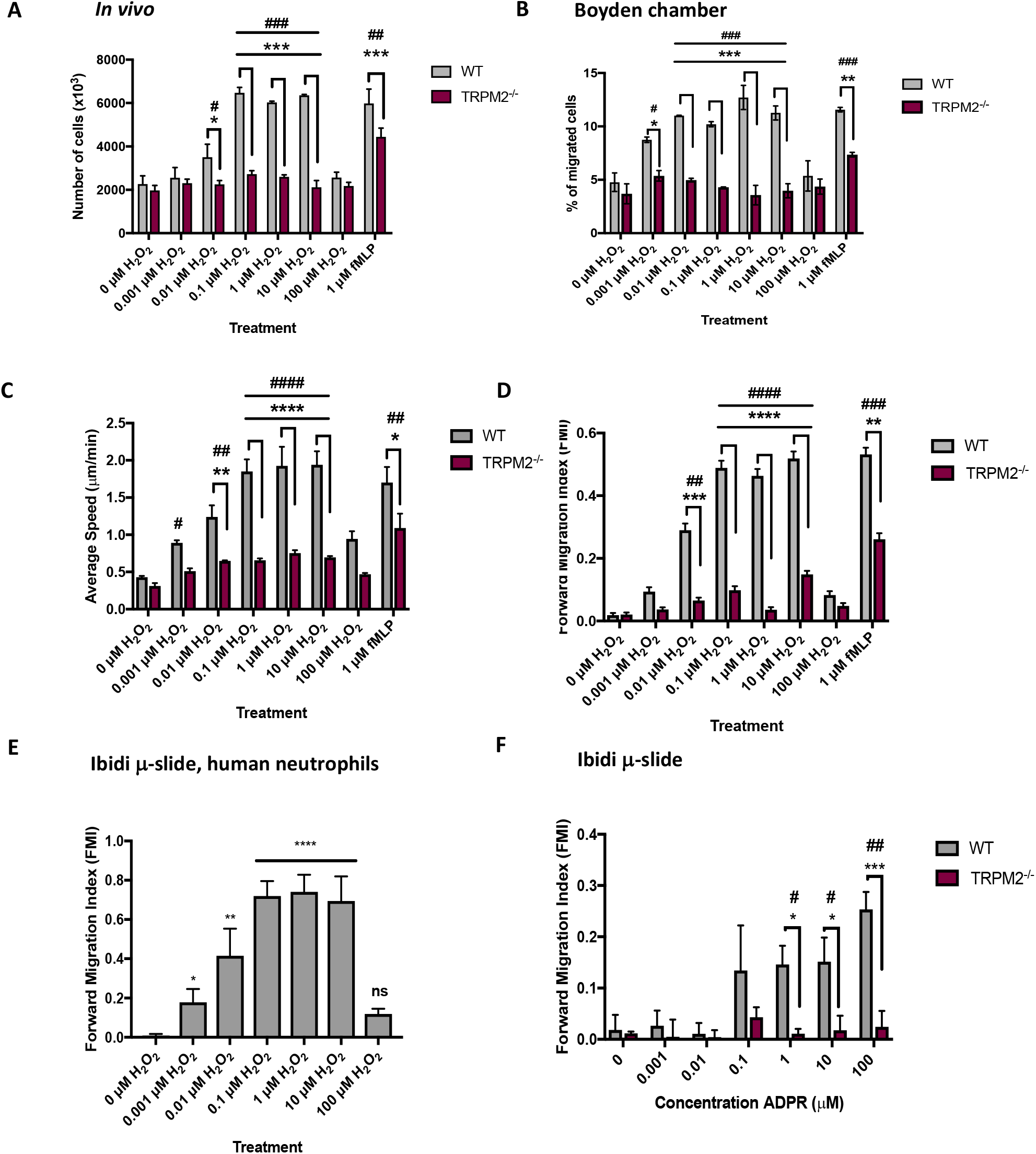
Neutrophil chemoattraction by H_2_O_2_ depends on TRPM2. A. Chemotaxis of neutrophils into the peritonea of WT and TRPM2^-/-^ mice following i.p. injection of H_2_O_2_. Direct injection of 0 – 100 μM H_2_O_2_ or 1 μM fMLP into the peritonea of WT and TRPM2^-/-^ mice (see Methods for details) elicited movement of total number of cells per peritoneum (ordinate). Time course of chemotaxis was determined in WT mice as shown in Supplementary Materials Fig. 1. Maximal chemotaxis in WT mice calculated from the average of 3 points between 60 and 120min. For TRPM2^-/-^ mice, chemotaxis was measured only at the 90min point. Each bar shows mean ± SEM (n=3 mice). B. Boyden chamber assay of mouse neutrophil chemotaxis. Each bar shows mean ± SEM % migration from n = 4 mice (n = 8 for 0 μM). C. *In vitro* ibidi μ-slide chemotaxis chamber assay using mouse neutrophils. Average speed of neutrophil migration, calculated for each neutrophil as path length of distance travelled divided by time. Each bar shows mean ± SEM of n = 4 mice. D. *In vitro* ibidi μ-slide chemotaxis chamber assay using mouse neutrophils. Forward migration index (FMI) is distance migrated in direction of gradient of H_2_O_2_ or fMLP divided by total distance travelled, from same experiments as shown in C. Note that concentrations of H_2_O_2_ in the nanomolar range are able to increase both speed and directionality of neutrophil migration, and that migration is inhibited at high concentrations of H_2_O_2_ (>10μM). Distribution of values of FMI shown in Supplementary Materials Fig. 2. Each bar shows mean ± SEM of n = 4 mice. E. *In vitro* ibidi μ-slide chemotaxis chamber assay using neutrophils isolated from human volunteer blood. FMI of human blood neutrophils shows similar dependence on concentration of H_2_O_2_ to mouse peritoneal neutrophils (compare with D). Each bar shows mean ± SEM of n = 3 individual human neutrophil samples. F. Extracellular ADPR is a potent chemoattractant dependent on TRPM2 activation. WT mouse neutrophils showed significantly increased FMI compared to TRPM2^-/-^ neutrophils in response to 1 – 100 μM ADPR. Each bar shows mean ± SEM from n = 3 mice. Statistical analysis: For comparison with 0 H_2_O_2_: #, p<0.05, ##, p<0.01, ###, p<0.001, ####, p<0.0001 (One-way ANOVA and Tukey-Kramer post-hoc test). For comparison with TRPM2^-/-^ (except in E): *, p<0.05, **, p<0.01, ***, p<0.001, ****, p<0.0001 (Two-way ANOVA and Bonferroni’s post-hoc test).

We next used the classical Boyden chamber method to measure neutrophil recruitment by H_2_O_2_ *in vitro.* In this assay, a membrane with 3 μm pores, through which only neutrophils can penetrate, separates the neutrophil and chemoattractant compartments. H_2_O_2_ down to 1nM concentration caused significant neutrophil recruitment through the membrane, with maximal recruitment between 10nM and 10μM H_2_O_2_ and a similar inhibition to that observed *in vivo* at 100μM (Fig. 1B).

Similar results were obtained using a second *in vitro* method, the ibidi μ-slide, in which neutrophil motion is continuously tracked under a microscope (Fig. 1C, D and Supplementary Video 1). Fig. 1C shows the speed of movement of neutrophils as a function of concentration of H_2_O_2_. Neutrophils undergo random movements in the absence of H_2_O_2_, so the average speed is not zero in the absence of H_2_O_2_ even though there is no significant directed movement (see Fig. 1D). The average speed is increased in a gradient of 1nM H_2_O_2_ distributed over 1mm, the distance between the compartments containing H_2_O_2_ and DMEM. Average speed is maximal between 0.1 μM and 10 μM H_2_O_2_ and is inhibited at 100μM H_2_O_2_. Maximal speed in fMLP is not significantly different from that in H_2_O_2_.

An alternative method of quantifying neutrophil movement is to measure the forward migration index (FMI), a measure of directionality, that gives the ratio of linear distance travelled in the direction of the H_2_O_2_ gradient to total distance travelled (Fig. 1D). A significant increase in FMI was observed at 10nM H_2_O_2_, and FMI was maximal between 0.1 μM and 10μM H_2_O_2_. As in the other assays, directed migration was inhibited at 100μM H_2_O_2_.

All three independent measures agree in showing that H_2_O_2_ is a highly potent neutrophil chemoattractant, from low nanomolar levels up to 10μM, and that chemoattraction is inhibited at concentrations of H_2_O_2_ above 10μM. A similar concentration-dependence of directed migration in a gradient of H_2_O_2_ was also observed with human blood neutrophils (Fig. 1E).

### Sensitivity of neutrophils to single molecules of H_2_O_2_

The evidence above shows that neutrophils exhibit a clear increase in both movement and direction-finding in concentration gradients as low as 10nM H_2_O_2_, distributed over the 1mm gap between the chemoattractant and DMEM compartments of the ibidi μ-slide. This gradient corresponds to a concentration difference between the leading and trailing edge of a 10μm diameter neutrophil of 100pM, and, assuming that H_2_O_2_ equilibrates rapidly and completely between the extracellular and intracellular spaces, this is a difference on average of around five intracellular molecules of H_2_O_2_ between the leading and trailing halves of the neutrophil (see calculation in legend to Supplementary Materials Fig. 2). The most sensitive neutrophils are therefore able to detect and respond reliably to only a few molecules difference of H_2_O_2_ between the leading and trailing edges. This extraordinary sensitivity to H_2_O_2_ suggests that the detection of H_2_O_2_ is a highly-evolved property, critical for survival.

**Fig. 2:**
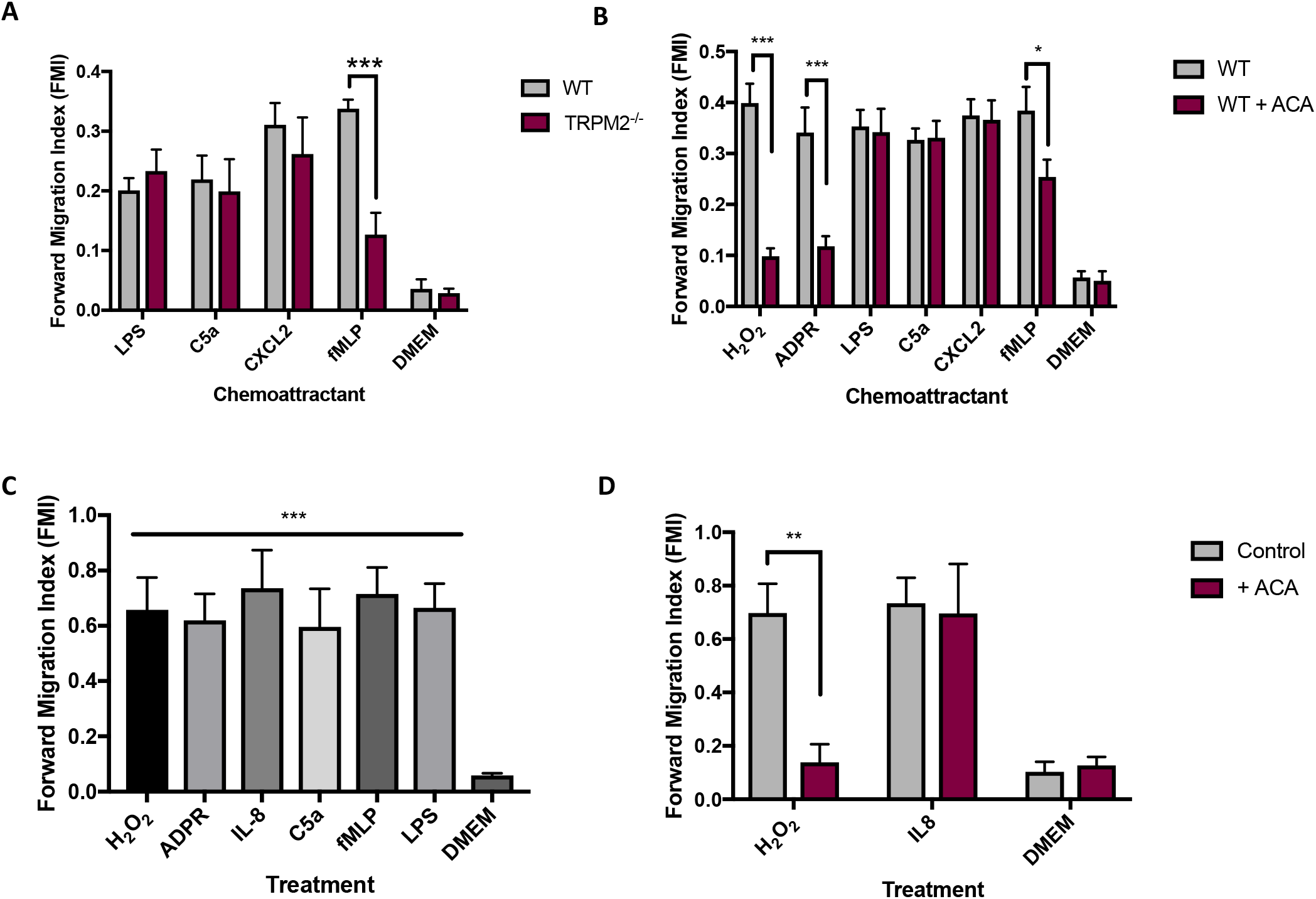
Dependence on TRPM2 of migration towards neutrophil chemoattractants. A. Effect of genetic deletion of TRPM2 on mouse neutrophil migration. Forward migration index (FMI) in response to gradient of lipopolysaccharide (LPS, 50 ng/ml), C5a (10 nM), CXCL2 (10 nM) and fMLP (1 μM), all over 1mm. No significant difference in FMI of WT and TRPM2^-/-^ neutrophils in response to LPS, C5a or CXCL2, but neutrophil migration to fMLP was significantly inhibited by TRPM2 deletion. B. Effect of pharmacological block of TRPM2 with ACA (N-(p-amylcinnamoyl)anthranilic acid, 10 μM) on mouse neutrophil migration. FMI in gradient of H_2_O_2_ (0.01 μM), ADPR (100 μM) and other chemoattractants as in A. C. Human blood neutrophil FMI in response to gradients of chemoattractants as follows, all over 1mm: H_2_O_2_ (0.01 μM), ADPR (100 μM), IL8 (10nM), C5a (10 nM), fMLP (1 μM) and LPS (50 ng/ml). D. Inhibition of human blood neutrophil FMI by TRPM2 blocker ACA (10 μM). Gradients over 1mm: H_2_O_2_ (0.01 μM), IL8 (10nM). Statistical analysis: Mean ± SEM, independent samples from n = 3 mice or humans in each panel. A, B, D: pairwise t-test comparing WT to TRPM2^-/-^ or ACA. C: One-way ANOVA and Tukey-Kramer post-hoc test compared to DMEM. *, p<0.05; **, p<0.01; *** p<0.001; ****, p<0.0001, difference non-significant if not stated.

### Neutrophil chemotaxis towards H_2_O_2_ depends on TRPM2

The TRPM2 ion channel is expressed in neutrophils and is potently activated by H_2_O_2_^12^, suggesting that it may be a candidate for the sensor of H_2_O_2_. TRPM2^-/-^ mice are highly susceptible to infection^13^, suggesting that the absence of TRPM2 causes a critical immune system defect. We found that genetic deletion of TRPM2 strongly inhibited H_2_O_2_-dependent neutrophil migration in all three assays (Fig. 1A-D), supporting the idea that TRPM2 is the major physiological sensor of H_2_O_2_ in neutrophils.

### Neutrophil chemotaxis towards ADPR depends on TRPM2

The intracellular C terminus of TRPM2 contains a nudix hydrolase homology domain (NUDT9-H), named after the mitochondrial ADP-ribose (ADPR) pyrophosphatase NUDT9, to which ADPR binds to activate the ion channel^14–21^. ADPR is known to cross the cell membrane, though the mechanisms are not clear^21,22^ (see also Fig. 3G below). We found that neutrophils responded vigorously to an extracellular gradient of ADPR (Fig. 1F). Migration up a gradient of ADPR was significantly diminished by deletion of TRPM2 (Fig. 1F), showing that neutrophil migration up a gradient of ADPR also depends on TRPM2.

**Fig. 3.**
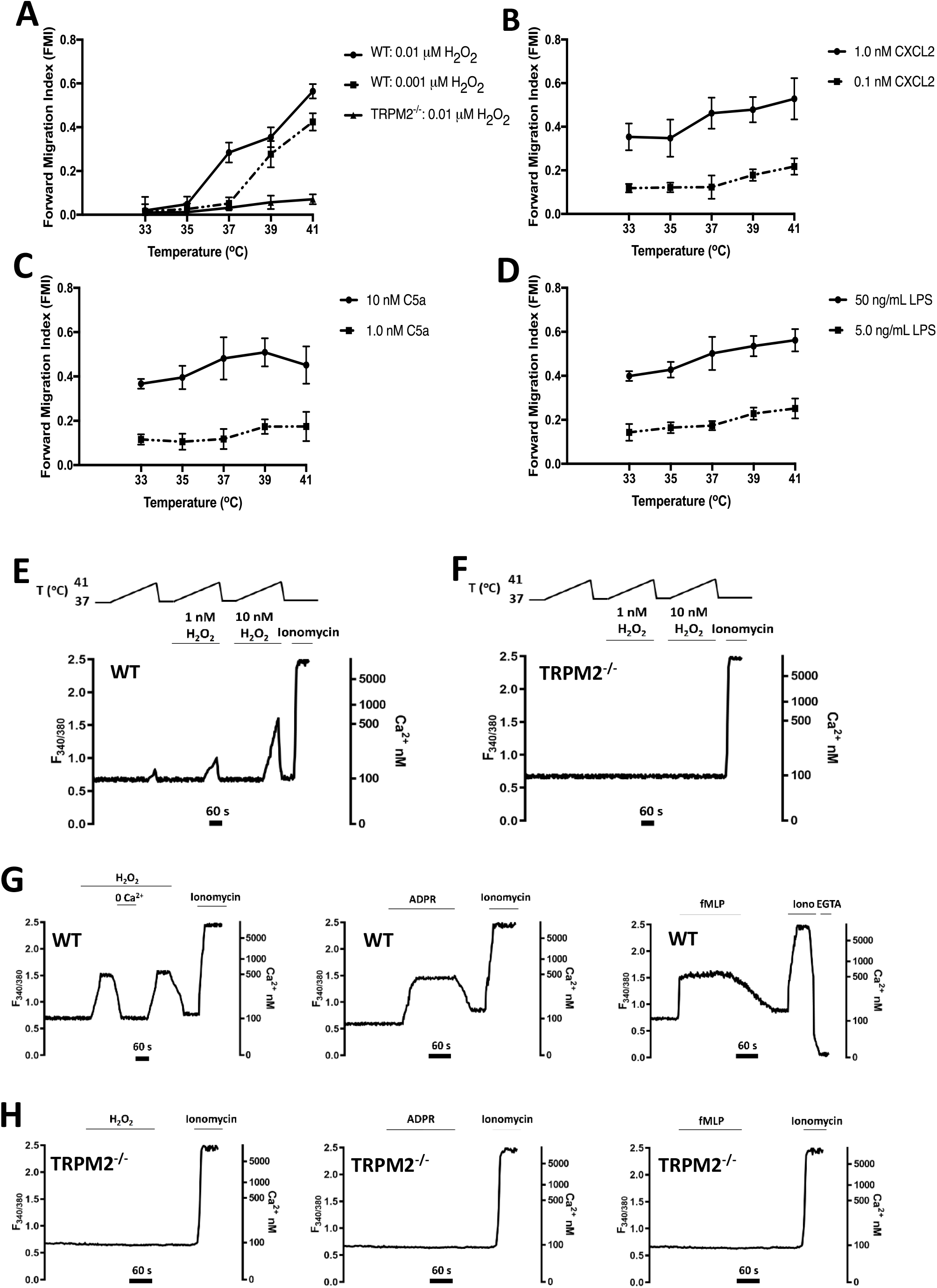
A-D: Dependence on temperature and TRPM2 of forward migration index (FMI) of neutrophil chemotaxis. Neutrophil migration in response to H_2_O_2_ tracked for 2 h in the ibidi μ-slide chemotaxis system using live-cell time-lapse microscopy. FMI, as a function of temperature, measured with the following genotypes and chemoattractants: (A) WT, to 0.001 μM and 0.01 μM H_2_O_2_; TRPM2^-/-^, to 0.01 μM H_2_O_2_; (B) WT, to 0.1nM and 1nM CXCL2; (C) WT, to 1.0nM and 10nM C5a; (D) WT, to 5.0 and 50ng/ml LPS. Concentrations of conventional chemoattractants giving sub-maximal and near-maximal migration determined in separate experiments (see Supplementary Materials Fig. 3). Each point shows mean ±SEM, n = 3 separate experiments using neutrophils from 3 mice. **E, F: Potentiation of calcium entry through TRPM2 by temperature and H_2_O_2_.** (E) Increase of [Ca]_i_ in response to a temperature ramp from 37°C to 41°C in neutrophils from WT mice. Threshold for [Ca]_i_ increase was 40.2°C in the absence of H_2_O_2_, 39.1 °C in 1nM H_2_O_2_ and 38.8°C in 10 nM H_2_O_2_. (F) No increase of [Ca]_i_ in neutrophils from TRPM2^-/-^ mice. Traces are from single neutrophils representative of 15-20 cells total in n=3 separate experiments using neutrophils from 3 mice; all traces had closely similar form. Neutrophil calcium imaging performed in flowing solution in absence of chemotactic gradient (see Methods). Calibration for conversion of F_340/380_ ratio to [Ca]_i_ shown in last panel in G (see Methods). **G, H: Calcium entry elicited by TRPM2-dependent chemoattractants H_2_O_2_, ADPR and fMLP.** (G) Increase of [Ca]_i_ in response to application of (from left) H_2_O_2_ (10μM), ADPR (100μM) and fMLP (1μM), in neutrophils from WT mice. In the H_2_O_2_ experiment (first panel) the [Ca]_i_ rapidly decreased when external calcium was removed (nominal 0Ca, no EGTA) consistent with Ca entry from the external solution via TRPM2 (Note contracted time scale in this panel). The trace in fMLP shows an example of the method used to convert F_340/380_ to [Ca]_i_ by measuring F_max_ with exposure to the Ca ionophore ionomycin (10μM), followed by F_min_ in 0Ca/2mM EGTA. (H) No increase of [Ca]_i_ in neutrophils from TRPM2^-/-^ mice. Experimental details as in E, F. All traces from single neutrophils but representative of 15-20 traces from n=3 separate experiments using 3 mice.

### Chemotaxis towards most conventional chemoattractants does not depend on TRPM2

The marked inhibition of neutrophil migration up gradients of H_2_O_2_ or ADPR, caused by deletion of TRPM2, could be due to an effect of the deletion on migratory ability *per se.* This possibility is not, however, supported by the observation that TRPM2 deletion does not affect chemotaxis to the known chemoattractants lipopolysaccharide (LPS), C5a and CXCL2 (Fig. 2A). Deletion of TRPM2, however, reduces but does not completely abolish chemotaxis towards the bacterial/mitochondrial peptide fMLP (Fig. 2A), as also noted in all assays shown in Fig. 1 A-D, suggesting that fMLP exerts its chemotactic effect in part by activating TRPM2.

### Neutrophil chemotaxis towards H_2_O_2_ or ADPR is inhibited by blocking TRPM2

To test further the dependence of neutrophil migration on TRPM2, we examined the effect of pharmacological block of TRPM2 with the inhibitor ACA (N-(p-amylcinnamoyl)anthranilic acid^23^). Block of TRPM2 with ACA caused a significant inhibition of chemotaxis in response to both H_2_O_2_ and ADPR (Fig. 2B and Supplementary Materials Fig. 1B), similar in magnitude to the effect of genetic deletion of TRPM2 shown in Fig. 1. Block of TRPM2 with ACA had no effect, however, on chemotaxis towards LPS, C5a and CXCL2, but caused a partial inhibition of migration towards fMLP (Fig. 2B). ACA has off-target actions^23^, but the close resemblance to results obtained with genetic deletion of TRPM2 supports the hypothesis that the effects of both genetic deletion and pharmacological block of TRPM2 are due to inhibition of TRPM2 itself and not to an off-target effect on another protein.

Fig. 2C shows similar experiments on human blood neutrophils. All chemoattractants strongly increased FMI to a similar extent, as was also observed with mouse peritoneal neutrophils (Fig. 2A, B). In Fig. 2D, block of TRPM2 with ACA in human blood neutrophils largely inhibited migration up a gradient of H_2_O_2_ but had no effect on migration up a gradient of IL-8, the human homologue of CXCL2, results closely resembling those in mouse neutrophils (Fig. 2B).

In summary, the results in Fig. 1 and 2 strongly support the hypothesis that TRPM2 is the cellular target that mediates guidance of neutrophils up a gradient of H_2_O_2_.

### Variability in values of FMI

There is a range of sensitivities to H_2_O_2_ amongst neutrophils, with some showing strongly directional migration up a gradient of H_2_O_2_ (FMI close to 1.0), while others show lower, or even absent, directional motility (Supplementary Materials Fig. 2 and Supplementary video 1). The variability in values for FMI amongst individual neutrophils explains why the mean FMI does not typically exceed a value of around 0.6, when a value of 1.0 would be expected if all neutrophils migrated in a straight line. The proportion of highly mobile to less mobile neutrophils was found to be variable in different experiments (Supplementary Materials Fig. 2). The reason for this diversity in response is unknown but could be related to the age or condition of different neutrophils in the pool.

We tested whether the apparently lower maximal values of FMI in some experiments (for example with ADPR, Fig. 1F) was due to a real difference or simply reflected variability between different neutrophil samples. Fig. 2A-C show that maximal FMI values are in fact not statistically different amongst H_2_O_2_, ADPR, LPS, C5a CXCL2 and fMLP when measured on a single neutrophil sample.

### Neutrophil migration to H_2_O_2_ depends strongly on temperature

TRPM2 is a member of the large TRP family of ion channels, and like several other members of this family, is strongly activated by increasing temperatures within the physiological range^20,24–26^. Fig. 3A shows that neutrophil migration towards very low concentrations of H_2_O_2_ is potently enhanced by small increases of temperature and that the effect of temperature is abolished by deletion of TRPM2. The effect of temperature is particularly prominent in a gradient of 1nM H_2_O_2_, when an elevation from 37°C to 39°C, corresponding to a mild fever, enhances the ability of neutrophils to migrate up a gradient of H_2_O_2_ from an FMI of 0.052 ± 0.028 to 0.277 ± 0.059, a 5.3-fold increase. By contrast, the effect of temperature on migration towards the conventional chemoattractants CXCL2, C5a and LPS is much less marked (Fig. 3 B-D). The strong temperature dependence of neutrophil migration towards H_2_O_2_ provides further support for a critical role of TRPM2 in neutrophil migration in response to H_2_O_2_, and moreover suggests a reason why fever may be beneficial in fighting infection, because an elevation of temperature by only 2°C, corresponding to a mild fever, gives a five-fold increase in neutrophil migratory ability towards low levels of H_2_O_2_.

### Temperature and chemoattractants cause elevations in intracellular calcium

TRPM2 is permeable to calcium ions^12^, like other members of the TRP family^27,28^, suggesting that the striking dependence of neutrophil migration on temperature may be due to a calcium influx through activated TRPM2 ion channels. Fig. 3E shows that elevations of temperature in the physiological range (from 37°C to 41 °C) induce an intracellular calcium increase, and that low concentrations of H_2_O_2_ (1nM and 10nM) enhance the amplitude of the calcium increase and lower its temperature threshold. Deletion of TRPM2 abolishes the thermally-induced elevations of calcium (Fig. 3F). The strong temperature-dependence of calcium influx through TRPM2 supports the hypothesis that a calcium influx through TRPM2 underlies the striking temperature-sensitivity of neutrophil migration driven by H_2_O_2_ (Fig. 3A). The results also support the idea that calcium influx through TRPM2 may be a driver of neutrophil motion, for which further evidence is presented below.

We next examined whether the agonists H_2_O_2_, ADPR and fMLP, whose effect on neutrophil migration depends on activation of TRPM2 (see above), also induce a calcium influx. Fig. 3G and H show that in all cases these agonists cause an elevation of intracellular calcium, from the resting level of around 100nM to 500nM, and that the increase is dependent on TRPM2. There was a significant delay between application of H_2_O_2_ or ADPR and activation of calcium influx, consistent with an intracellular site of action for both agonists. H_2_O_2_ is able to cross the cell membrane through aquaporins^29^, and ADPR is also known to cross the neutrophil cell membrane, though the mechanisms are not clear^21,22^. There was a rapid drop in intracellular calcium when external calcium was removed (Fig. 3G, left-hand panel), showing that activation of TRPM2 causes an influx of calcium from the extracellular solution.

### Leading-edge calcium pulses in H_2_O_2_ and ADPR are driven by TRPM2

The results above suggest that preferential activation of TRPM2 channels by elevated levels of H_2_O_2_ at the neutrophil leading edge could cause a localised calcium influx that guides neutrophil chemotaxis by coupling to the motile machinery. We used calcium imaging of migrating neutrophils and we observed that an elevated intracellular calcium concentration at the neutrophil leading edge is indeed seen during movement of a neutrophil up a gradient of H_2_O_2_ (Fig. 4A, left panel). Leading-edge intracellular calcium elevations were seen in all migrating neutrophils, and they had a pulsatile appearance for which the name “calcium pulse” seemed appropriate (see for example Supplementary video 2). The calcium pulse is most prominent at the base of the pseudopodium, but in some images it can be seen also invading the advancing pseudopodium itself (Fig. 4B, C and Fig. 5A, B). Genetic deletion of TRPM2 abolishes both calcium pulses and cell migration (Fig. 4A, centre, and Fig. 4B - E). At high concentrations of H_2_O_2_, maximal activation of TRPM2 in all parts of the cell membrane means that a uniformly high level of calcium floods the cell and neutrophil migration is abolished (Fig. 4A, right).

**Fig. 4.**
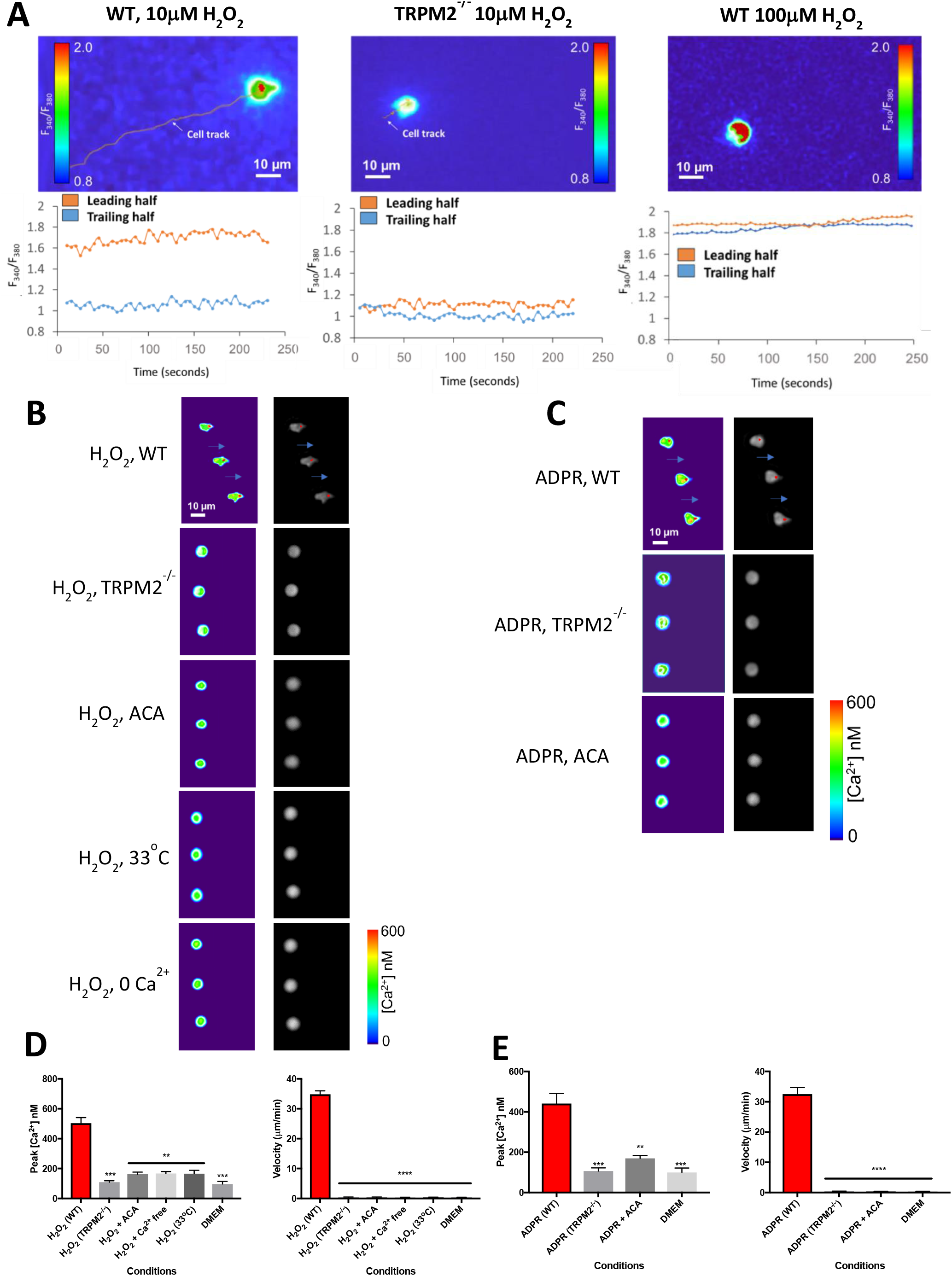
Neutrophil chemotaxis is accompanied by leading-edge calcium pulses. A. Images show calcium-dependent fluorescence ratio of indicator fura2 with alternating 340nm and 380nm illumination. Left: Wild-type neutrophils migrating up a gradient of H_2_O_2_ (10μM over 1mm, left to right) show elevated intracellular calcium at the leading half of the cell in the direction of the gradient (see lower graph). Centre: When TRPM2 is deleted, neutrophils show no net migration, and the difference between leading and trailing edge calcium is close to zero. Right: In a higher concentration gradient of H_2_O_2_ (100μM) migration is inhibited and calcium concentration is uniformly high throughout the cell. Images representative of n =3 individual experiments. Similar experiments in Supplementary Fig. 4 show effect of ADPR and temperature. B. Chemotaxis towards is H_2_O_2_ accompanied by leading-edge calcium pulses. Left: colour-coded calcium images (scale shown at bottom). Right: maximal calcium concentration (red) superimposed on brightfield phase-contrast images. Both chemotaxis and calcium pulses are abolished by deletion or pharmacological block of TRPM2 with ACA (10μM), by withdrawal of extracellular calcium or by lowering the temperature to 33°C. Fast-moving individual neutrophils selected for imaging. Images acquired at 0.5s intervals; time between each image shown is 10s. Long blue arrows indicate movement of neutrophil by a distance equal to or above the cell diameter in the time interval of 10s. Gradient of H_2_O_2_ 10 μM over 1mm, except bottom panel which is 10nM over 1mm. Other concentrations as in Fig. 2. Images representative of n =3 individual experiments. C. Calcium pulses and chemotaxis in a gradient of ADPR (100 μM) are abolished by genetic deletion of TRPM2 and by the TRPM2 blocker ACA (10μM). Other details as in B. D. Peak calcium (left) and cell velocity (right) for the conditions shown in B. Peak [Ca^2+^] averaged over n = 40 images in each series (sample images shown in B). Velocity calculated as in Fig. 1C. Bars show average of n = 3 individual cells. One-way ANOVA and Tukey-Kramer post-hoc test compared to first bar. *, p<0.05; **, p<0.01; ***, p<0.001; ****, p<0.0001 E. Similar plots for data shown in C.

**Fig. 5.**
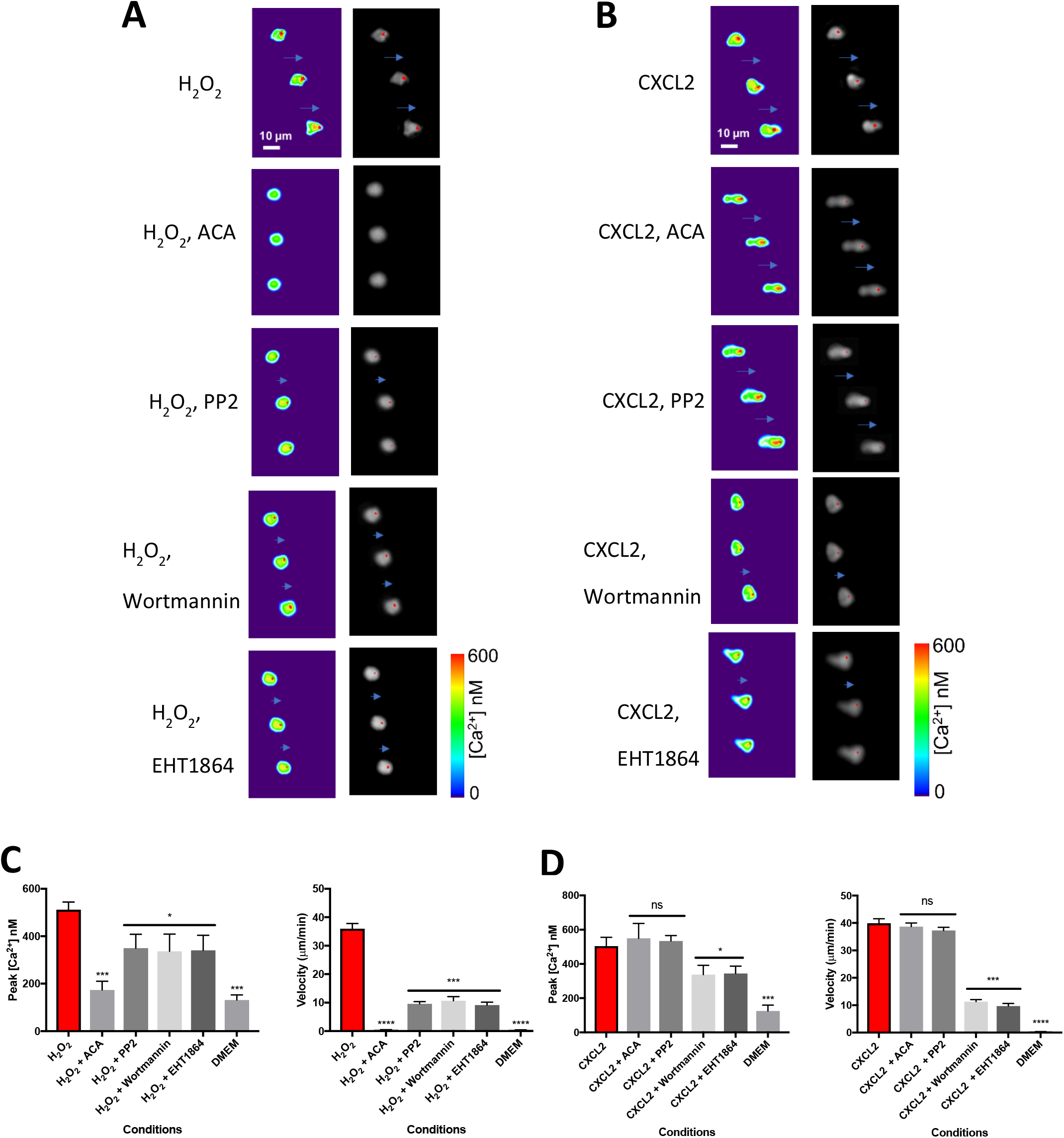
Effect of pharmacological blockers on neutrophil migration towards H_2_O_2_ and chemokine CXCL2. A. Gradient of H_2_O_2_ (10μM over 1mm). From top: H_2_O_2_ alone; TRPM2 blocker ACA (N-(p-amylcinnamoyl)anthranilic acid, 10μM) completely inhibits both migration and Ca pulses (reproduced from Fig. 4); migration is slowed and Ca^2+^ pulses partially inhibited by the Src-family kinase inhibitor PP2 (10 μM), PI3kinase inhibitor wortmannin (100 nM) and Rac inhibitor EHT 1864 (50 μM). Images shown at 10s intervals in each panel. Long blue arrows - movement of neutrophil a distance equal to or greater than the cell diameter; arrowheads - movement less than cell diameter; no arrow – no significant movement. Other details as in Fig. 4. Images representative of n =3 individual experiments. B. Similar experiment with gradient of CXCL2 (10 nM). Migration and Ca^2+^ pulses are unaffected by ACA (10 μM) and PP2(10 μM), while migration is slowed and Ca^2+^ pulse amplitude reduced by wortmannin (100 nM) and ETH 1864 (50 μM). Images representative of n =3 individual experiments. C. Peak calcium (top) and cell velocity (bottom) for the conditions shown in A. Bars show average of n = 3 individual cells. Two-way ANOVA and Tukey-Kramer post-hoc test compared to first bar. *, p<0.05; ***, p<0.001; ****, p<0.0001, ns = not significant. Other details as in Fig. 4. D. Similar plots for data shown in B. Similar experiments for conventional chemoattractants C5a and LPS shown in Supplementary Materials Fig 5.

Pharmacological block of TRPM2 with the inhibitor ACA^30^ has a similar effect to genetic deletion of TRPM2 in inhibiting calcium pulses and neutrophil chemotaxis towards H_2_O_2_ and ADPR (Fig. 4B - E). Both calcium pulses and neutrophil movement are also abolished by withdrawal of extracellular calcium (Fig. 4B, D), consistent with an influx of calcium from the external medium acting as a trigger for the generation of intracellular calcium pulses. Lowering the temperature to 33°C, which strongly inhibits neutrophil motility (Fig. 3A), also inhibits the generation of calcium pulses (Fig. 4B, D and Supplementary Materials Fig. 4B). These parallel observations using a range of approaches support the idea that leading-edge calcium pulses in both H_2_O_2_ and ADPR are generated by an influx of calcium through TRPM2, and that calcium pulses are coupled to migration.

### Leading-edge calcium pulses in classical chemoattractants are not driven by TRPM2

Fig. 5B and Supplementary Materials Fig. 5 show that leading-edge calcium pulses, similar to those observed in H_2_O_2_ and ADPR, are also seen in gradients of the chemoattractants CXCL2, LPS and C5a. These observations suggest that calcium pulses “steering” neutrophil chemotaxis, by determining the future direction of establishment of pseudopodia, may be a universal mechanism. Block of TRPM2 with ACA has no effect on calcium pulses in the classical chemoattractants CXCL2, LPS and C5a. Together with the data in Fig. 2 showing the lack of effect of TRPM2 deletion or block on neutrophil migration to classical chemoattractants, these experiments show that neither the generation of leading-edge calcium pulses nor neutrophil migration in these chemoattractants is driven by a TRPM2-dependent mechanism.

### Signalling pathways determining neutrophil motility

There have been several proposals regarding the mechanisms that couple chemoattractants to the intracellular motile machinery of immune cells, including activation of PI3K to generate PIP3 at the neutrophil leading edge, leading to activation of Rac^1,31–34^; or, in the case of chemoattraction by H_2_O_2_, oxidative modulation of cysteine residues present in the Src family tyrosine kinase *lyn*^9,11^. Fig. 5 and Supplementary Materials Figs. 5 and 6 investigate potential signalling pathways involved in neutrophil chemotaxis. PP2, a potent inhibitor of Src family kinases (SFK), caused only partial inhibition of motility in a gradient of H_2_O_2_ (Fig. 5A, C and Supplementary Materials Fig. 6) suggesting that a member of the Src kinase family modulates H_2_O_2_-directed motility but is not essential for generating movement. PP2 also partially inhibited the amplitude of calcium pulses (Fig. 5A, C), consistent with a SFK member acting upstream of the generation of calcium pulses. Interestingly, PP2 had no effect on either neutrophil motility nor generation of calcium pulses in gradients of the classical chemoattractants CXCL2, LPS or C5a (Fig. 5B, D and Supplementary Materials Fig. 5, 6), showing that the effect of SFK inhibition is specific to the H_2_O_2_/TRPM2 mechanism. One possibility is that *lyn* may promote trafficking of TRPM2 to the surface membrane, and thus inhibition of *lyn* would reduce the calcium influx triggered when TRPM2 is activated by H_2_O_2_. An SFK member has been shown to play a similar role in promoting trafficking of TRPV1 to the neuronal cell surface membrane^35^.

**Fig. 6.**
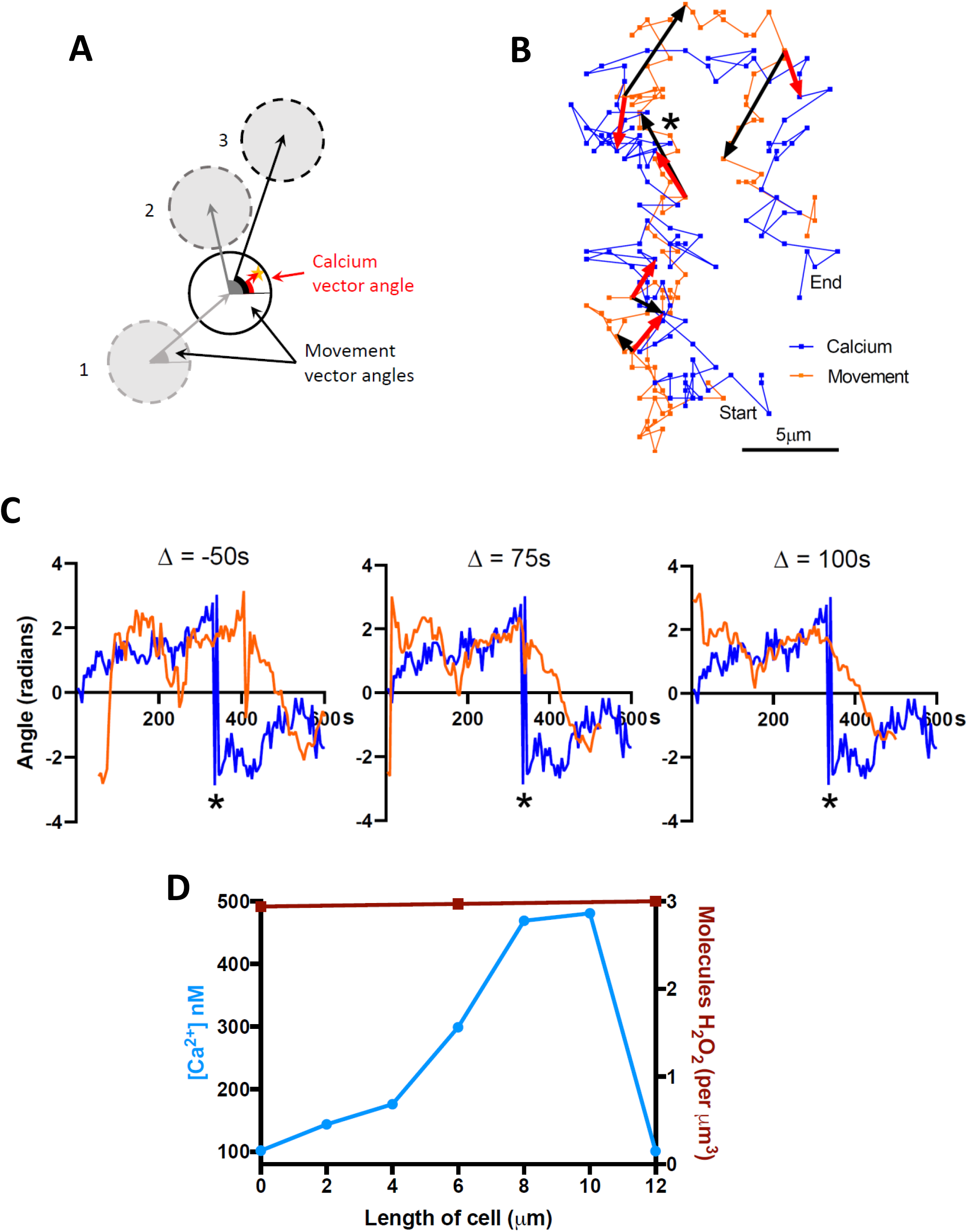
Calcium pulses direct neutrophil chemotaxis. A. Method for determining the relation between direction of calcium pulses and direction of neutrophil movement. Red vector (“calcium vector”) shows direction in which the calcium pulse is “pointing”, by connecting cell geometric centre to simultaneous location of maximal calcium signal, while grey or black vectors show direction of movement of cell geometric centre (“movement vector”), which can be calculated at earlier (1) or later times (2, 3). B. Tracks of geometric cell centre (orange) and position of peak calcium concentration (blue) at 5s intervals for a neutrophil moving in absence of chemotactic gradient. (See movie in Supplementary Video File 2). Similar image series taken at a 10x shorter intervals did not show any obvious difference in the dynamics of movement of peak calcium concentration (Supplementary Materials Fig. 8). Red vectors (shown at 100s intervals) give direction of calcium vector (see A). Black vectors show movement vector (see A) connecting the geometric centre at the same time as each calcium vector with the geometric centre 50s later. At the time indicated by the star the calcium vector shifts abruptly from an average upward orientation to an average downward orientation. There is a clear delay before the movement vector follows. C. Calcium vector (blue) and movement vector (orange) angles for cell shown in B. A delay of 75s best predicts the moment at which the movement vector begins to follow the abrupt change in direction of calcium vector. There is a further delay before the cell movement aligns with the new calcium vector orientation (half-time c. 83s), giving total delay between change of calcium vector and half-alignment of movement vector of 158s. D. Calcium concentration (blue), determined from fura-2 fluorescence ratio, shown in the in the direction of migration from left to right of a neutrophil located half-way up a gradient of H_2_O_2_ from 0 to 10nM. For comparison, red line shows H_2_O_2_ concentration (in molecules of H_2_O_2_ per μm^3^) along the length of the cell. Horizontal axis gives distance from left-hand to right-hand edges of the cell, in the direction both of the H_2_O_2_ gradient and of cell motion.

PI3K and Rac have also been suggested as initiators of neutrophil chemotaxis^1,31–34^. However, the PI3K inhibitor wortmannin and the Rac inhibitor EHT1864 only partially inhibited neutrophil chemotaxis in response to H_2_O_2_, CXCL2, LPS and C5a (Fig. 5 and Supplementary Materials Fig. 5, 6). In the case of chemotaxis to H_2_O_2_, the residual motility in the presence of the PI3K and Rac blockers was completely inhibited by the TRPM2 blocker ACA (Supplementary Materials Fig. 6A). The PI3K and Rac inhibitors also reduced the amplitudes of leading-edge calcium signals, suggesting that PI3K and Rac play a modulatory role in the pathways upstream of the generation of calcium pulses, but that the amplitude of calcium pulses is the primary event controlling motility in response to all chemoattractants.

### Calcium pulses determine the direction of neutrophil chemotaxis

Increases in intracellular calcium in neutrophils and other immune cells have been suggested to be a consequence of cell movement, as might arise if a calcium increase was required to retract the trailing edge of the cell^36^, or because an interaction between cell and substrate triggers calcium entry caused by the movement itself^37,38^. In these scenarios, the calcium increase would be expected to follow the movement, while if the calcium pulse directs the movement it would be expected to precede it. For this reason we investigated whether a change in the cellular location of calcium pulses precedes a subsequent change in the direction of cell migration (calcium drives migration), or whether a change in the direction of migration precedes a change in location of calcium pulses (migration drives calcium). Neutrophils in the absence of a chemotactic gradient still exhibit calcium pulses, which change cellular location at intervals, and also exhibit random migration (See Supplementary video 2), providing a useful way of distinguishing these two possibilities.

We quantified the angle between the geometric centre of the cell and the centre of the calcium pulse occurring at the same moment (calcium vector angle, Fig. 6A) and compared this angle with the angle between the geometric centre of a neutrophil at the moment of the calcium pulse and at different times as the cell advances (movement vector angle, Fig. 6A). In the case of the neutrophil shown in Fig. 6B and Supplementary Video File 2, the calcium vector turns from an overall upward direction to an overall downward direction halfway through the recording (shown by a star in Fig. 6B and 6C). As is also visible in Supplementary Video File 2, the change in cell direction follows the change in calcium signal direction with a significant delay.

The calcium vector connecting the cell geometric centre and the calcium pulse is measured at a given moment (blue trace in Fig. 6C), but the movement vector (orange trace) can refer to the cell position at a variable past or future time. Fig. 6C shows that a lag of +75s between calcium vector and movement vector best describes the moment at which the movement vector begins to reverse in response to the abrupt change in the direction of the calcium vector. This positive delay is consistent with calcium driving the direction of movement, while a negative delay gives a poor fit (shown for a −50s delay). These results are consistent with a change in calcium pulse location driving a change in movement, with a delay of around 75s, and does not agree with the idea that movement might cause (i.e. anticipate) the calcium pulse. There is a further significant delay (half-time ~83s) before the cell movement fully aligns with the new direction of the calcium vector (see middle panel in Fig. 6C), giving a total time lag between change of the direction of the calcium pulse and half-time of alignment of movement with the new calcium pulse location of 158s. Two further neutrophils analysed in the same way gave similar delays, with total half-time for the alignment of cell movement to a change in calcium direction of 146s and 164s. The neutrophil movement therefore reacts to a change in the direction of a calcium pulse with a surprisingly long delay, with total half-time of around 160s, or almost 3 minutes.

Finally, Fig. 6D shows a comparison between the external H_2_O_2_ concentration surrounding a cell migrating half-way up a 10nM gradient over 1mm, and the internal calcium concentration measured with fura-2. The calcium gradient is very much steeper than the H_2_O_2_ gradient, showing that there must be a powerful non-linear amplification process that converts the gentle H_2_O_2_ gradient into the strongly enhanced calcium signal seen at the cell leading edge.

## Discussion

The work described here confirms previous studies showing that H_2_O_2_ is a potent neutrophil chemoattractant both *in vitro*^2^ and *in vivo*^3^. We demonstrate an essential role for the TRPM2 ion channel in directing neutrophil chemotaxis towards H_2_O_2_. We propose that preferential activation of TRPM2 by the higher levels of H_2_O_2_ at the neutrophil leading edge causes a calcium influx, triggering amplified leading-edge calcium pulses that determine the direction of extension of pseudopodia and therefore of future neutrophil motion. There is a significant delay, of around 160s, between a change of location of a calcium pulse and the half-time of response of the neutrophil direction, in striking contrast to the millisecond reaction time of many other calcium-driven cellular processes such as skeletal muscle contraction or synaptic vesicle exocytosis.

We find that neutrophils are guided by H_2_O_2_ over four orders of magnitude of concentration, from 1nM to 10μM. Higher concentrations of H_2_O_2_ (>10μM) inhibit chemotaxis and cause an elevated calcium concentration throughout the cell. In agreement, previous studies have also found inhibition of chemotaxis in neutrophils and lymphocytes at levels of H_2_O_2_ above 10μM^39,40^. A probable cause of the inhibition of chemotaxis at higher H_2_O_2_ is loss of the differential intracellular calcium gradient necessary for cell guidance, as shown in Fig. 4A.

Neutrophil guidance towards conventional chemoattractants such as LPS, C5a or CXCL2 is independent of TRPM2, but similar leading-edge calcium pulses are still observed, suggesting that guidance by calcium pulses is a universal mechanism. Signalling from the G-protein coupled receptors activated by conventional chemoattractants to intracellular calcium stores is a likely mechanism^41,42^. An interesting exception to the TRPM2-independence of conventional chemoattractants is chemotaxis towards fMLP, which is partially inhibited by either pharmacological block or genetic deletion of TRPM2. A possible explanation may lie in the recently-discovered physical interaction between TRPM2 and FPR1, the receptor for fMLP^40^.

Previous studies of the involvement of TRPM2 in neutrophil chemotaxis have come to contradictory conclusions. Yamamoto et al^43^ found that deletion of TRPM2 decreased neutrophil migration in response to an inflammatory supernatant formed by activated neutrophils *in vitro* and also decreased movement to areas of inflammation *in vivo.* In support, Hiroi et al^44^ found that neutrophil invasion following myocardial ischaemia was lowered in Trpm2^-/-^ mice. Wang et al^40^, in contrast, found that deletion of TRPM2 *promotes* chemotaxis both *in vitro* and *in vivo* and they propose that TRPM2 generates a “stop” signal to halt neutrophil migration in the vicinity of pathogens. In the present study we show both in vitro and in vivo that neutrophil motility is potently promoted by low levels of H_2_O_2_, and that motility is inhibited, both *in vivo* and *in vitro*, by either pharmacological inhibition or genetic deletion of TRPM2. Thus, at low concentrations of H_2_O_2_, TRPM2 mediates a powerful chemoattractant mechanism. In addition, we find that high concentrations of H_2_O_2_ (above 10μM) inhibit neutrophil motility, which may be critical in providing a “stop” mechanism in the vicinity of pathogens. We show that inhibition is due to a rise of the internal calcium level throughout the cell when TRPM2 channels are maximally activated by H_2_O_2_, thus flooding the cell with calcium and abolishing the internal calcium gradient necessary for neutrophil navigation.

The ratio between calcium at the leading and trailing edge of an advancing neutrophil is much greater than the ratio between H_2_O_2_ concentrations (Fig. 6D), suggesting that a process of non-linear amplification enhances the leading-edge calcium signal. One likely possibility for such a mechanism is the highly non-linear process of calcium-induced calcium release from subcellular stores^21,22,41,42,45^.

A scheme consistent with the work reported here is shown in Supplementary Materials Fig. 8. In a gradient of H_2_O_2_, preferential activation of TRPM2 ion channels at the leading edge causes an influx of calcium from the extracellular solution, triggering calcium-induced calcium release from intracellular stores. The Src family kinase *lyn* modulates this pathway at an early stage, perhaps by regulating trafficking of TRPM2 ion channels to the surface membrane. The intracellular signalling molecules PI3K and Rac, which have been proposed as drivers of chemotaxis, do not appear to play a direct role in chemotaxis, but instead fine-tune motility to all chemoattractants by regulating a step common to all chemoattractants, for example refilling of the intracellular calcium stores, and thus modulate the amplitude of the leading-edge calcium pulses.

Some studies have found no evidence for calcium gradients in neutrophils during chemotaxis^38,46^, though others have obtained evidence that leading-edge calcium signals play a critical role in chemotaxis of many different cell types^47,48^. In the present work, we observed leading-edge calcium signals in all motile neutrophils *in vitro.* Interestingly, leading-edge calcium signals have also recently been detected in migrating neutrophils *in vivo*^49^. These authors find that neutrophils can be guided *in vivo* by expressing TRPV1 and then imposing a directional gradient of the TRPV1 agonist capsaicin^49,50^. This work supports the idea that a TRP channel alone can mediate neutrophil guidance, but it does not identify the responsible TRP isoform. Here we identify TRPM2 as the ion channel mediating physiological neutrophil guidance towards a source of H_2_O_2_. TRPM2 may be an interesting future drug target, for instance in controlling excess neutrophil invasion in conditions such as sepsis.

Neutrophils are able to navigate up a gradient of H_2_O_2_ in which there is an average difference of only a few molecules of intracellular H_2_O_2_ between the leading and trailing halves of the cell. This extraordinary sensitivity suggests that navigation towards H_2_O_2_ is vital for mammalian survival and has been highly honed by evolution. The strong temperature dependence of TRPM2 activation^20,24–26^ suggests a novel function for fever: by enhancing TRPM2 activation, fever potently enhances the sensitivity to a gradient of H_2_O_2_, thereby enhancing neutrophil guidance and promoting the detection and killing of invading pathogens. H_2_O_2_ may therefore play a more critical role than conventional cytokines in the early phase of the response of the innate immune system to tissue damage or pathogen attack.

## Materials and Methods

### Animals

Black C57BL/6 WT and TRPM2^-/-^ K/O mice (6-8 weeks old) were bred in house from matings of TRPM2^+/-^ heterozygous mice to ensure that the genetic and environmental backgrounds of both WT and TRPM2-/- mice were as far as possible identical. TRPM2^-/-^ K/O mice were a gift from the Mori laboratory (Kyoto, Japan)^43^ and had been backcrossed onto the WT C57BL/6 strain for nine generations. For experiments in which comparisons with K/O were not required, Black C57BL/6 WT mice (6-8 weeks old) were purchased from Charles River Inc. All animal work was conducted under UK Home Office personal and project licences, approved by the Animal Welfare Ethical Review Board (AWERB) of King’s College London and carried out in accordance with The Animals (Scientific Procedures) Act 1986 and in compliance with the ARRIVE guidelines.

### Chemicals and reagents

Hydrogen peroxide (H_2_O_2_, 31642), adenosine 5’diphosphoribose (ADPR, A0752), *N*-formylmethionine-leucyl-phenylalanine (fMLP, F3506), thioglycolate (70157), lipopolysaccharide (LPS, LPS25), CXCL2 (SRP4251), complement component 5a (C5a, SRP4895A), *N*-(p-Amylcinnamoyl)anthranilic acid (ACA, A8486), PP2 (P0042), interleukin-8 (IL-8, CXCL8 SRP3098) were purchased from Sigma-Aldrich (Sigma-Aldrich Company Ltd., Dorset, UK). Wortmannin (W1628) was purchased from Yorlab (York, UK) and EHT1864 (3872/10) was purchased from Bio-Teche (Minnesota, US). Pluronic F-127 (P3000MP) and Fura-2AM (F1221) were purchased from Thermo Fisher Scientific (Thermo Fisher Scientific Life Technologies, Waltham, Massachusetts, U.S). The RAL DIFF-QUIK kit (a modified version of the May-Grünwald-Giemsa stain)) (RAL555) was purchased from RAL Diagnostics, (RAL Diagnostics, Martillac, France).

### *In vivo* model: measurement of H_2_O_2_-induced peritoneal cell chemotaxis

An *in vivo* model was developed in which WT and TRPM2 K/O mice were subjected to intraperitoneal (IP) injection (10 μl/g) of: 0, 0.001, 0.01, 0.1, 1, 10 or 100 μM H_2_O_2_ or 1 μM fMLP in H_2_O. After a variable time mice were euthanised, skin covering the peritoneum was removed, 5 ml PBS injected into the peritoneal cavity and the cavity massaged gently for 60s to dislodge cells. The peritoneal fluid was extracted and centrifuged for 10 mins at 200 RCF. The supernatant was discarded and the cells suspended in DMEM (Thermo Fisher Scientific Cat. No. 41966-029) + 10% FBS. The total number of cells recovered was counted, using a Neubauer chamber, as a semi-quantitative measure of cell chemotaxis. Cell numbers were corrected for the amount of fluid recovered as a fraction of 5ml.

### Isolation of mouse peritoneal neutrophils

Mice were injected IP with 3% thioglycolate solution (10 μl/g) and, after 4 h, euthanised by cervical dislocation. The peritoneal-covering skin was removed, 5 ml PBS injected into the peritoneal cavity and massaged gently for 60 s to dislodge cells. The peritoneal fluid was extracted and centrifuged for 10 mins at 200 RCF. The supernatant was discarded and cells suspended in DMEM + 10% FBS. This method generated cell suspensions containing >90% of neutrophils, identified through a fast-acting modified version of the May-Grünwald-Giemsa staining^51^ and subsequent cell type identification (see Supplementary Materials Figure 9).

### Cell identification in peritoneal extract

Cell suspension isolated from peritonea of WT mice as above. Suspensions spun down onto glass slides using a Cytospin at 400 RPM for 5 mins and left to air-dry overnight. A modified version of the May-Grünwald-Giemsa staining was used to identify cell types (RAL DIFF-QUIK kit, RAL diagnostics). Slides were suspended in RAL Diff-Quick fixative solution (methanol based solution to stabilize cellular components) for 1 minute, in RAL Diff-Quik solution I (Xanthene solution; a buffered solution of Eosin Y) for 1 minute and in RAL Diff-Quik solution II (a buffered solution of thiazine dyes, consisting of methylene blue and Azure A) for 1 minute. Nuclei were meta-chromatically stained red/purple and cytoplasm pink/yellow (See Supplementary Materials Fig. 9).

### Isolation of human blood neutrophils

Blood samples were collected from healthy volunteers by Dr Lucy Norling at Queen Mary College, University of London, with written consent. Human cells were prepared according to a protocol approved by the East London & The City Local Research Ethics Committee (no. 06/Q605/40; P/00/029 ELCHA, London, United Kingdom) and kindly donated for use in our experiments. Blood was collected into 3.2% sodium citrate and diluted 1:1 in RPMI 1640 before separation through a double-density gradient as described^52^.

### *In vitro* Boyden chamber neutrophil migration assay

QCM Boyden chamber chemotaxis neutrophil migration assay kits were purchased from Merck Millipore (ECM505, Massachusetts, USA). Neutrophil chemotaxis cell migration assays were conducted according to manufacturer’s instructions. In brief, neutrophils, isolated and prepared as above were seeded at 2×10^5^ cells per insert and each insert placed in a chamber containing either DMEM medium alone, 0.001 μM, 0.01 μM, 0.1 μM, 1 μM, 10 μM, 100 μM H_2_O_2_ or 1 μM fMLP. Inserts and chambers were incubated at 37°C in 95% air/5% CO_2_ for 4 hrs to allow cells to migrate through neutrophil-specific 3 μm pores. Non-migrated cells were removed from the inserts and the inserts placed in cell-detachment solution for 30 mins at 37°C in 95% air/5% CO_2_. 112 μl of this solution and of the lower chamber were added to the same wells of black-walled 96 well plates. Cells were lysed in the wells and incubated with CyQUANT GR dye, which emits increasing fluorescence proportional to the levels of bound nucleic acid released from the lysed cells. Following a 15 min incubation at room temperature, fluorescence intensity was analysed on a FLUOstar Omega microplate reader (BMG LABTECH, Buckinghamshire, UK) at 480/520 nm.

### Neutrophil chemotaxis assays

Ibidi μ-slide chemotaxis assay chambers, precoated with collagen IV along the central migration strip, were purchased from Thistle Scientific Ltd (Uddingston, Glasgow, UK). Neutrophils, isolated as above from peritonea of WT or TRPM2 K/O mice or from human blood, were re-suspended within 30min of collection in DMEM + 10% FBS at a concentration of 5×10^5^ cells per ml and 6μl was seeded along the central migration strip of an Ibidi μ-slide chamber as per the manufacturer’s instructions. Slides were incubated for 1 hr at 37°C in humidified 95% air/5% CO_2_, to allow neutrophil adherence to the central migration strip. DMEM (without added FBS) with and without added chemoattractant was then added to the wells on opposite sides of the central migration strip. DMEM was from Thermo Fisher Scientific Cat. No. 41966-029, or for experiments in which the effects of calcium removal were studied, 0Ca DMEM was from Thermo Fisher Scientific Cat. No. 21068-028. For experiments in which effects of inhibitors were to be tested, equal concentrations were added to both DMEM + chemoattractant and DMEM wells. Slides were pre-incubated at 37°C in 95% air/5% CO_2_ for 20 mins to allow the generation of a gradient of chemoattractant across the 1 mm wide x 70μm deep central cell migration strip. Live-cell time-lapse microscopy was then conducted using a 10x lens and dark-field illumination on a Nikon Eclipse Ti-E inverted microscope equipped with the Nikon Perfect Focus System (PFS). The images shown are centered on the middle of the 1mm wide cell migration strip and span the majority of its width. The microscope was housed in a temperature-controlled Perspex box (Solent Scientific) at 37°C, or other temperatures as desired, with slides housed in a stage-mounted block in humidified 95% air/5% CO_2_. A maximum of 12 individual chambers (4 individual slides, 3 chambers per slide) could be imaged per experiment by using a motorized stage. Stage movement, lens focus and image acquisition were controlled by Nikon NIS Elements software. Experiments were conducted over 2 h, with images of each assay compartment taken every 2 min. The ImageJ Fiji TrackMate plug-in was employed to track individual neutrophils. A chemotaxis and migration plug-in, provided by ibidi, was used to calculate speed and forward migration index (FMI) data from the neutrophil tracks.

### Calcium imaging of neutrophils

Neutrophils isolated as above were re-suspended in DMEM + 10% FBS at a concentration of 5×10^5^ per ml.

For experiments in which intracellular calcium was to be recorded while extracellular solutions were changed, neutrophils were plated onto a collagen-coated 13mm round glass coverslip and incubated at 37°C in 95%air/5% CO_2_ for 1h to allow neutrophils to adhere. Fura2-AM (5 uM in DMEM) was then added to the cells on the coverslip for 30 mins at 37°C in 95% air/5% CO_2_. Fluorescence was measured during alternating illumination at 340nm and 380 nm (OptoScan; Cairn Research Inc, Kent, UK) every 2 seconds using a Nikon Eclipse Ti inverted microscope with a 40x lens and iXon 897 EM-CCD camera controlled by WinFluor 3.2 software. F_340/380_ ratios were obtained using FIJI (ImageJ) and converted to calcium concentrations using the equation given by Grynkiewicz et al with values R_max_ = 2.501, R_min_ = 0.103, both determined experimentally, see Fig. 3G. Full details in Vilar et al^53^.

For experiments when calcium signals during chemotaxis up a gradient of chemoattractant were to be recorded, 1 μl of Fura-2 AM solution (50 μg Fura-2 AM + 10 μl pluronic F-127 + 10 μl DMSO) was added to 500 μl of neutrophil suspension and incubated for 1 hr at 37°C in 95%air/5% CO_2_. Fura-2 loaded cells in suspension were seeded into ibidi chambers as previously and imaged in a Nikon Ti-E microscope with a 40x phase contrast lens. Fast-moving neutrophils located in the middle of the central cell migration strip were selected, with typically only one cell imaged per field. Calcium ratio images were obtained with alternating 340nm and 380nm epi-illumination supplied by stable LED light sources (Fura-LED, Cairn Research), typically at 5s intervals. In some experiments acquisition of each pair of epifluorescence images was followed by a transmitted white-light phase-contrast image. All images were filtered by a broad-band 510nm filter and captured with a Photometrics Prime 95B sCMOS camera. Stage movement, focus and image acquisition were controlled by Nikon NIS Elements software. The ImageJ Fiji RatioPlus plug-in was used to generate F_340/380_ ratio images and a rainbow look-up table (LUT) was applied to the ratio images to indicate the level of calcium. Values of the ratio F_340/380_ were converted to intracellular free calcium concentration using the equation given by Grynkiewicz et al.^54^. To determine the values of fluorescence ratios at maximal (R_max_) and zero (R_min_) calcium levels, neutrophils were imaged in the presence of ionomycin (10μM) either in DMEM or in 0Ca DMEM with added 2mM EGTA, giving mean values R_max_ = 3.132 and R_min_ = 0.803 (note that the differences between these values and those above arises because chemotaxis experiments were performed on different apparatus). To indicate the location of the maximum calcium level on a phase-contrast image, a 3-colour LUT was applied to an F_340/380_ ratio image and the red channel (highest F_340/380_) was selected and superimposed on the phase-contrast image. The coordinates of the region of highest calcium within the cell were identified in F_340/380_ images using the Fiji maximum intensity identifying tool. The coordinates for the geometric centre of the neutrophil were identified using the cell edge and centre localising tool. In some images (for example, Fig. 4A) mean values of F_340/380_ in leading and trailing halves was calculated by identifying the cell outline using the cell edge localising tool and dividing the cell by hand into leading and trailing halves in the direction of the imposed chemotactic gradient.

## Acknowledgements

Supported by a grant from the Wellcome Trust to PMcN and by a KCL Biomedical Research Centre PhD studentship to HM. SL acknowledges the Department of Health Research (DHR), Ministry of Health & Family Welfare, Government of India for awarding a Long-Term Fellowship at KCL, UK. We thank Dr Larissa Pinto for assistance with histochemistry and Dr Lucy Norling and Professor Mauro Perretti for supplying human blood neutrophils.

