## Supplementary information for "TRPM2 ion channels steer neutrophils towards a source of hydrogen peroxide"

### Supplementary Materials Fig. 1

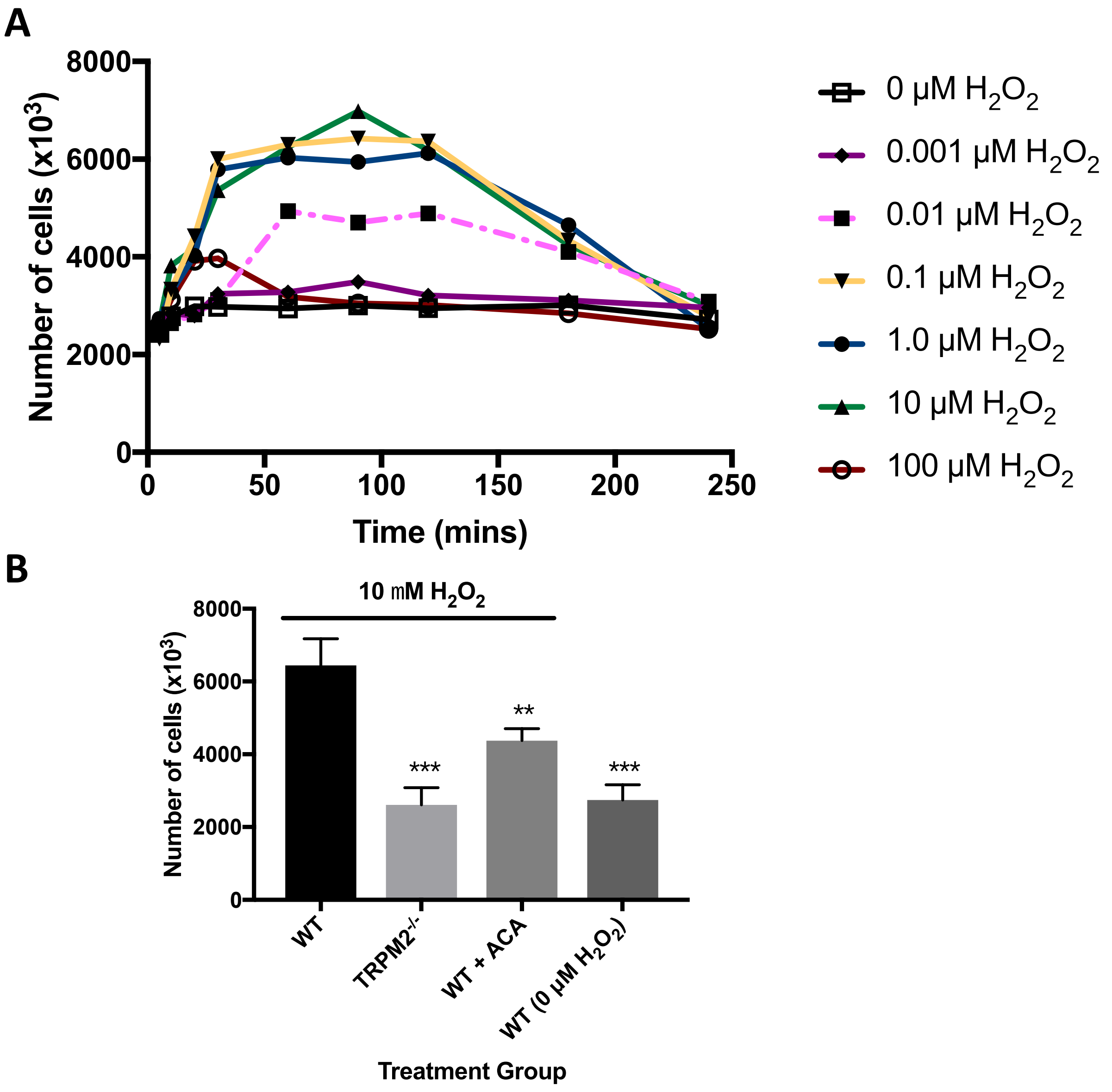


**Supplementary Materials Fig. 1. Chemotaxis of neutrophils into mouse peritoneum in response to i.p. injection of $H_2O_2$.**

A. Experiment performed as in Fig. 1A. After a variable time of incubation, mice were euthanised, cells were harvested from the peritonea and the recruitment of cells quantified by counting in a Neubauer chamber. More than 90% of harvested cells were neutrophils (see Methods). Recruitment reached a stable plateau between 60 and 120 min (black bar) and values at the times of 60, 90 and 120min were therefore averaged to determine maximum neutrophil movement (Fig. 1).
B. Genetic deletion of TRPM2 or i.p. injection of the TRPM2 blocker ACA (10 μM) both inhibit movement of neutrophils into the peritoneum. **, p<0.01; ***, p<0.001; (One-way ANOVA and Tukey-Kramer post-hoc test). Mean ± SEM, n = 3 mice in each panel.

### Supplementary Materials Fig. 2

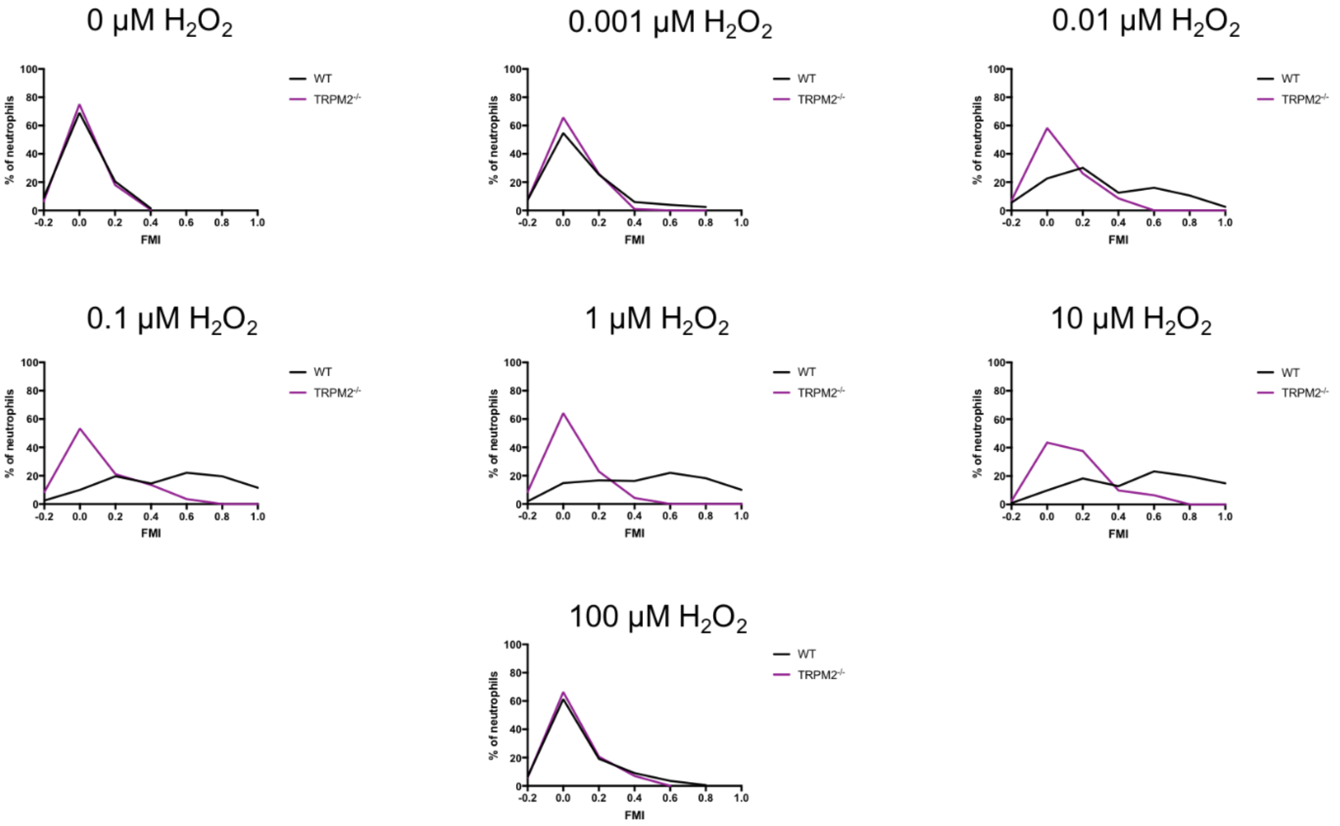


**Supplementary Materials Fig. 2. Distributions of FMI values for WT and TRPM2$^{-/-}$ neutrophils at increasing concentrations of $H_2O_2$.**

Data from Figure 1C (n =4) plotted as percentage of cells migrating with each value of FMI. WT, black; TRPM2$^{-/-}$, purple. The figure shows that neutrophils form two overlapping populations: a hypermobile population that responds strongly to $H_2O_2$ and a less-mobile population that responds more slowly or not at all to $H_2O_2$. Migration in response to different concentrations of $H_2O_2$, tracked for 2 hrs in the ibidi µ-slide chemotaxis system using live-cell time-lapse microscopy. Neutrophils isolated from mouse peritoneum as described in Methods.

**What is the difference between the number of intracellular molecules of $H_2O_2$ in the front and the back halves of a neutrophil navigating up a 10nM gradient of $H_2O_2$?**

Volume of a neutrophil represented approximately as a disc 10µM in diameter and 2µM thick is $\pi x(5)^2$ x $2\mu m^3 \approx 1.6 \times 10^{-13}$ litres. Neutrophils can detect and navigate up a linear gradient of 10nM over 1mm (see Fig. 1), so the threshold extracellular concentration difference between front and back is ≈100pM. Assuming rapid equilibration between the extracellular and intracellular concentrations and no degradation of intracellular $H_2O_2$, then the front half of a neutrophil contains $0.8 \times 10^{-23}$ moles more than the back or approximately 5 more molecules of $H_2O_2$.

### Supplementary Materials Fig. 3

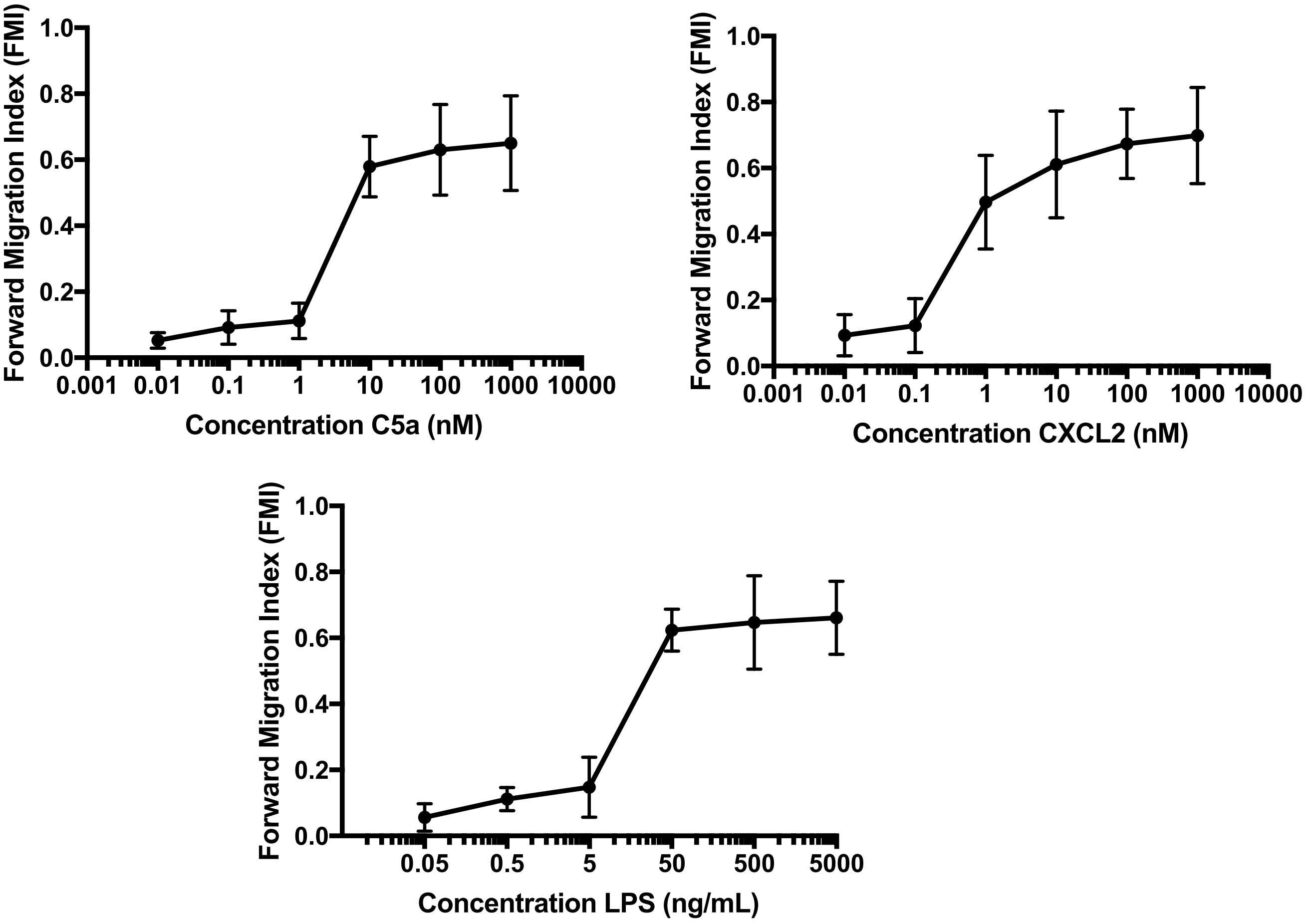


**Supplementary Materials Fig. 3. Neutrophil forward migration index (FMI) as a function of dose of the conventional chemoattractants C5a, CXCL2 and LPS.**

Experiment performed to determine concentrations giving below-maximal and near-maximal migration, determined from forward migration index (FMI), for use in experiments determining temperature-dependence of FMI in response to conventional chemoattractants (Fig. 3).

### Supplementary Materials Fig. 4

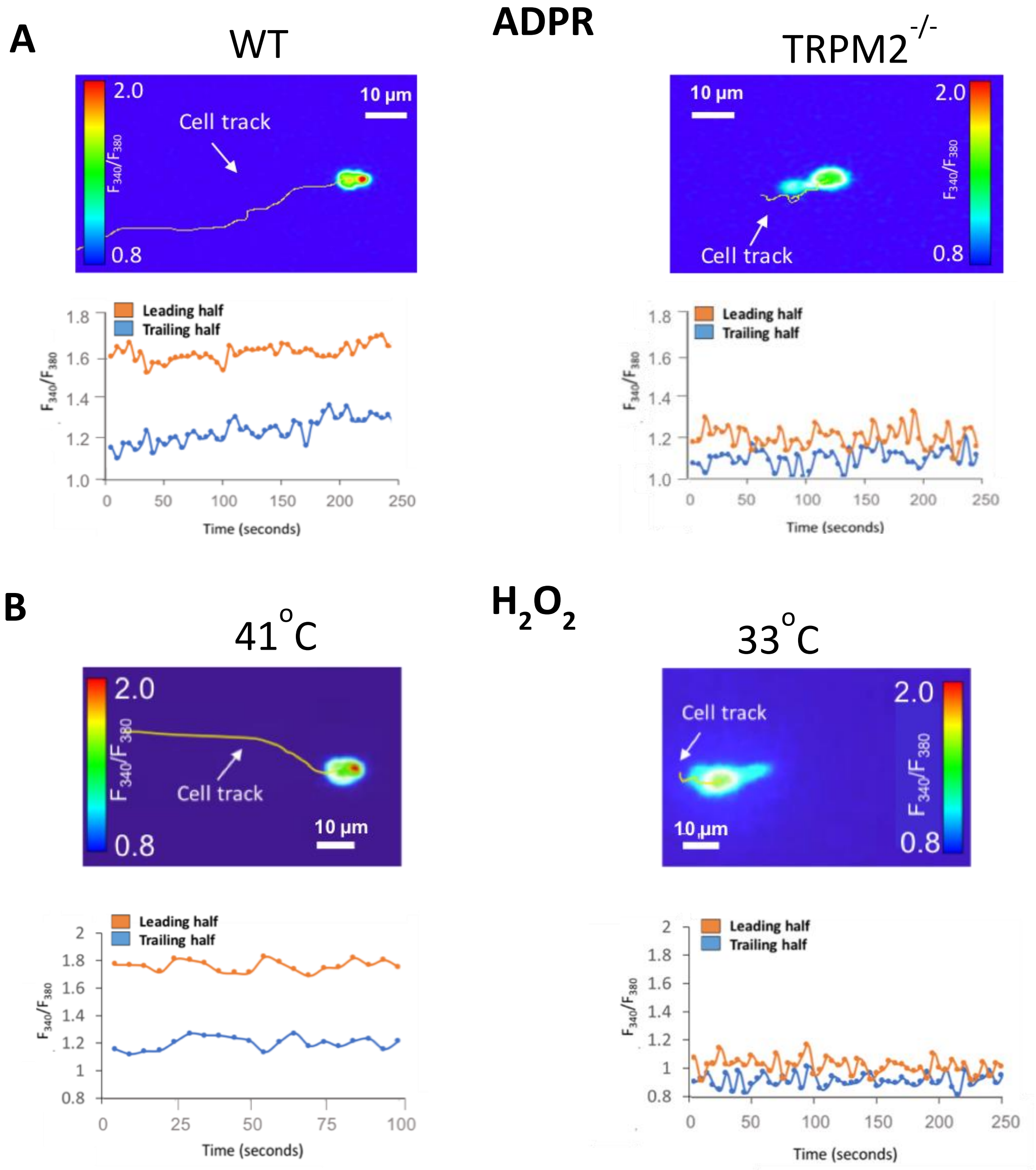


**Supplementary Information Fig. 4. Association between neutrophil motility and leading-edge calcium pulses: ADPR and temperature.**

A. Left: Images show calcium-dependent fluorescence ratio of indicator fura2 with alternating 340nm and 380nm illumination. Wild-type neutrophils migrating up a gradient of ADPR (10μM over 1mm, left to right) show elevated intracellular calcium at the leading edge (see lower graph). Right: When TRPM2 is deleted, neutrophils show no net migration, and the difference between leading and trailing edge calcium is close to zero. Images representative of n =3 individual experiments.

B. Similar experiment performed with neutrophils migrating up a gradient of $H_2O_2$ (10nM over 1mm, left to right) at 41°C (left) and 33°C (right). Images representative of n =3 individual experiments.

### Supplementary Materials Fig. 5

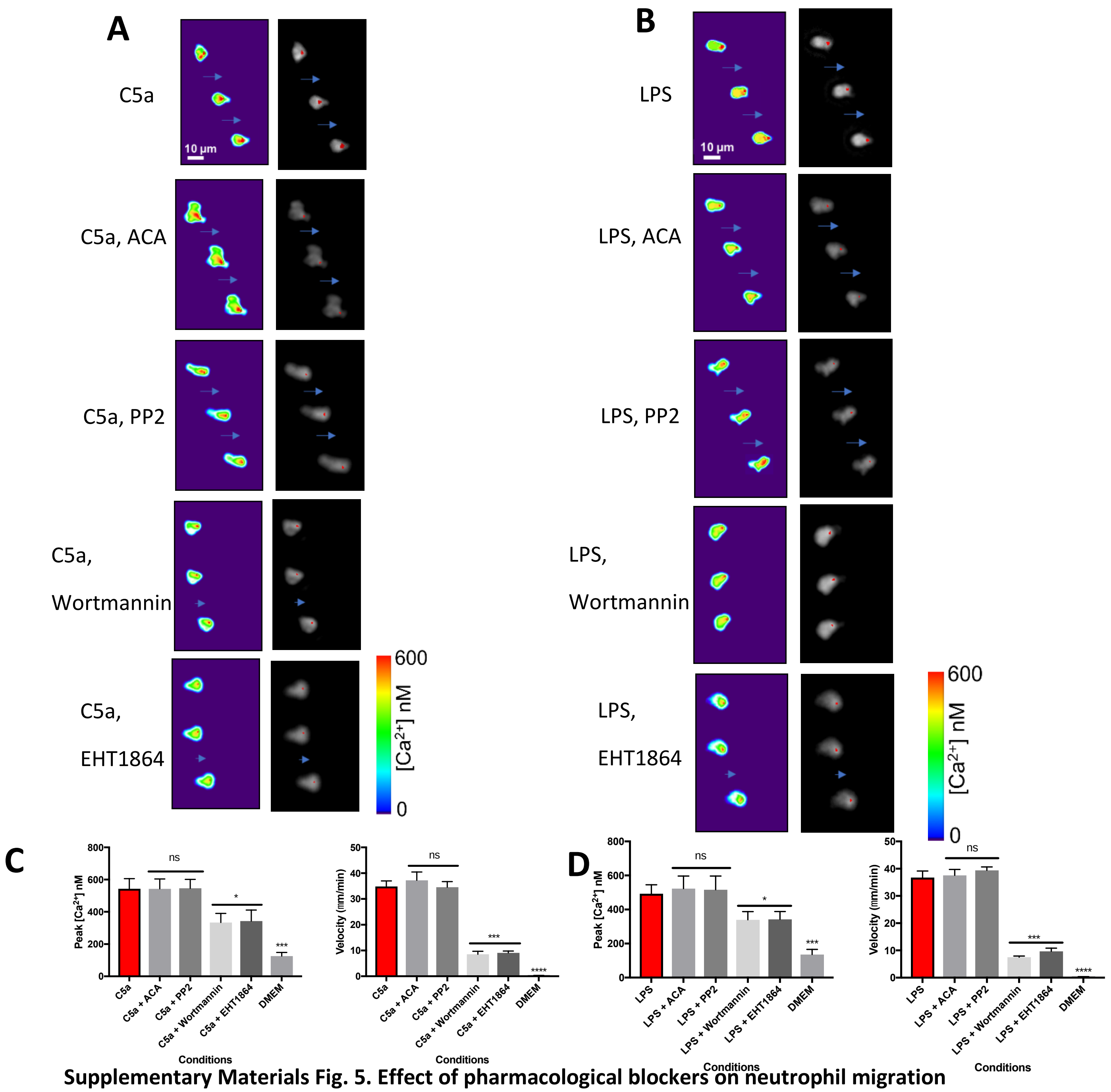


**Supplementary Materials Fig. 5. Effect of pharmacological blockers on neutrophil migration towards classical chemoattractants.**

A. Similar experiment to those shown in Fig. 4. Gradient of LPS (lipopolysaccharide, 50 ng/ml). Migration and Ca pulses are unaffected by ACA (10 µM) and PP2(10 µM), while migration is slowed and Ca pulse amplitude reduced by wortmannin (100 nM) and ETH 1864 (50 µM). Other details as in Fig. 4.
B. Similar experiment with gradient of C5a (10 nM).
C. Peak $[Ca^{2+}]$ and cell velocity in gradient of C5a, calculated as in Fig 4. ns = not significant; *, p<0.05; ***, p<0.001; ****, p<0.0001 (One-way ANOVA and Tukey-Kramer post-hoc test). Mean ± SEM, n = 3 mice in each panel.
D. Peak $[Ca^{2+}]$ and cell velocity in gradient of LPS (details as in C).

### Supplementary Materials Fig. 6

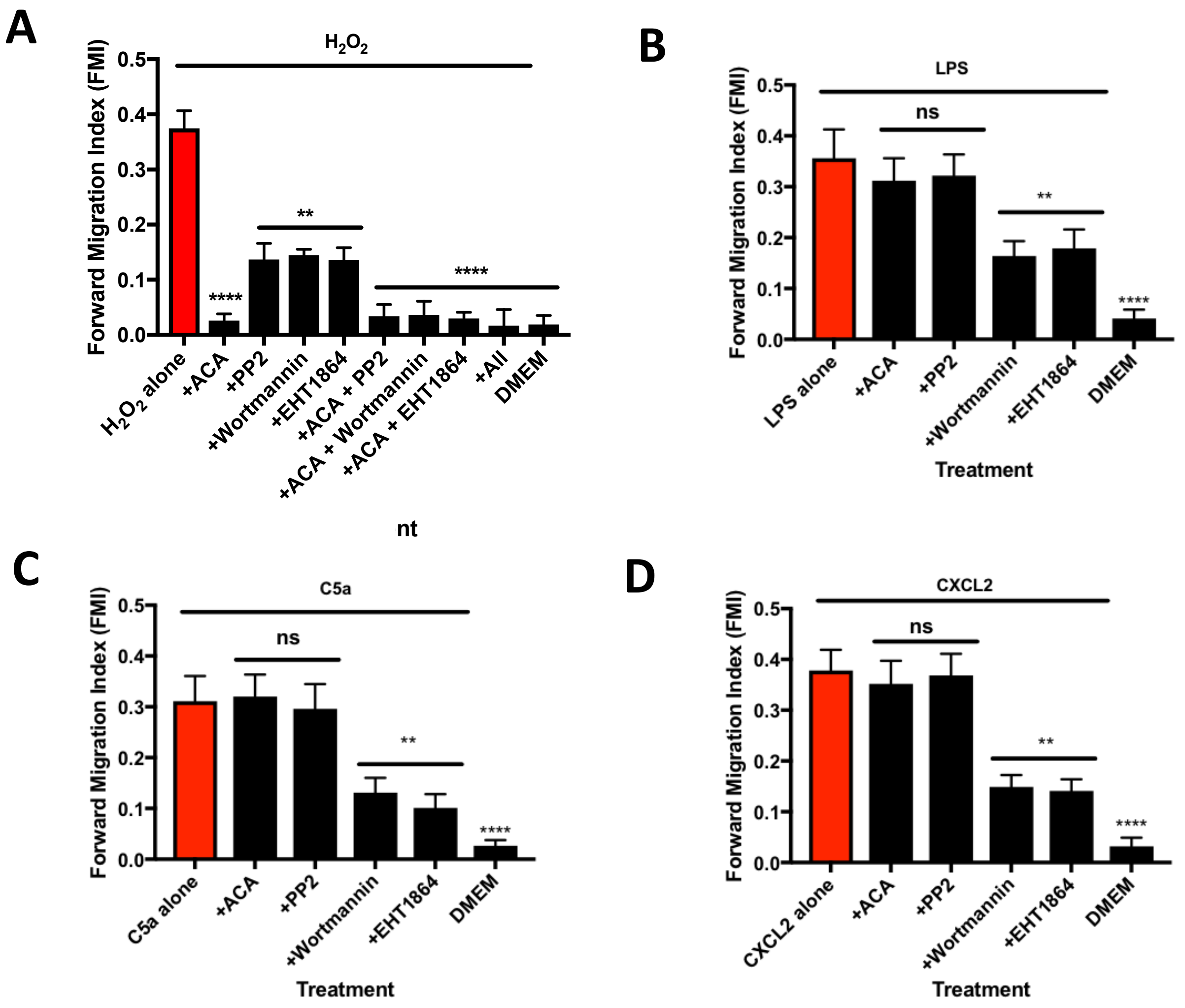


**Supplementary Materials Figure 6. Signalling pathways involved in neutrophil chemotaxis.**

A: Inhibition of neutrophil migration to 10nM $H_2O_2$ by TRPM2 inhibitor ACA (10 µM), Src family kinase (SFK) inhibitor PP2 (10 µM), PI3K inhibitor Wortmannin (100 nM) and Rac2 inhibitor EHT1864 (50 µM) all significantly reduced neutrophil chemotaxis. Neutrophils tracked as in Fig. 1, without calcium imaging.

B – D: Similar experiments with lipopolysaccharide (LPS, 50 ng/ml), C5a (10nM) and CXCL2 (10nM).

Statistics: Comparison with chemoattractant alone: ns, not significant; **, p<0.01;; ****, p<0.0001 (One-way ANOVA and Tukey-Kramer post-hoc test). Mean ± SEM, n = 3 mice in each panel.

### Supplementary Materials Fig. 7

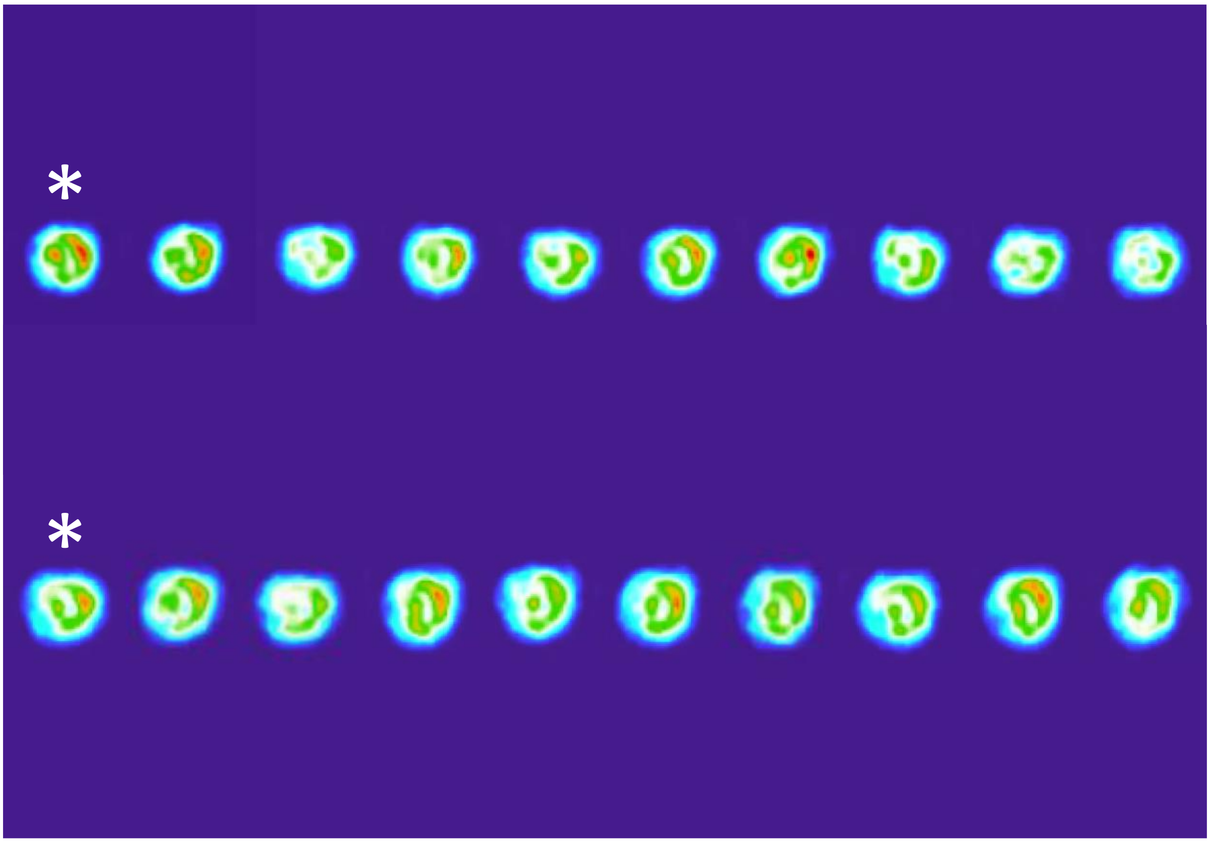


**Supplementary Materials Figure 7. Images of cell moving in absence of gradient taken every 500ms.**

Examples of images of neutrophil moving freely from left to right in absence of chemotactic gradient, with Ca images taken every 500ms. Images that would be sampled every 5s, as in Fig. 6, are indicated by stars. The distribution and amplitude of calcium elevations within the cell is to some extent variable but calcium concentration is highest at the leading edge in all images and there is no major change when images are taken at the slower image acquisition rate shown by the stars. Images have been displaced for clarity and distances between images do not represent actual movement in the 500ms interval between each image.

### Supplementary Materials Fig. 8

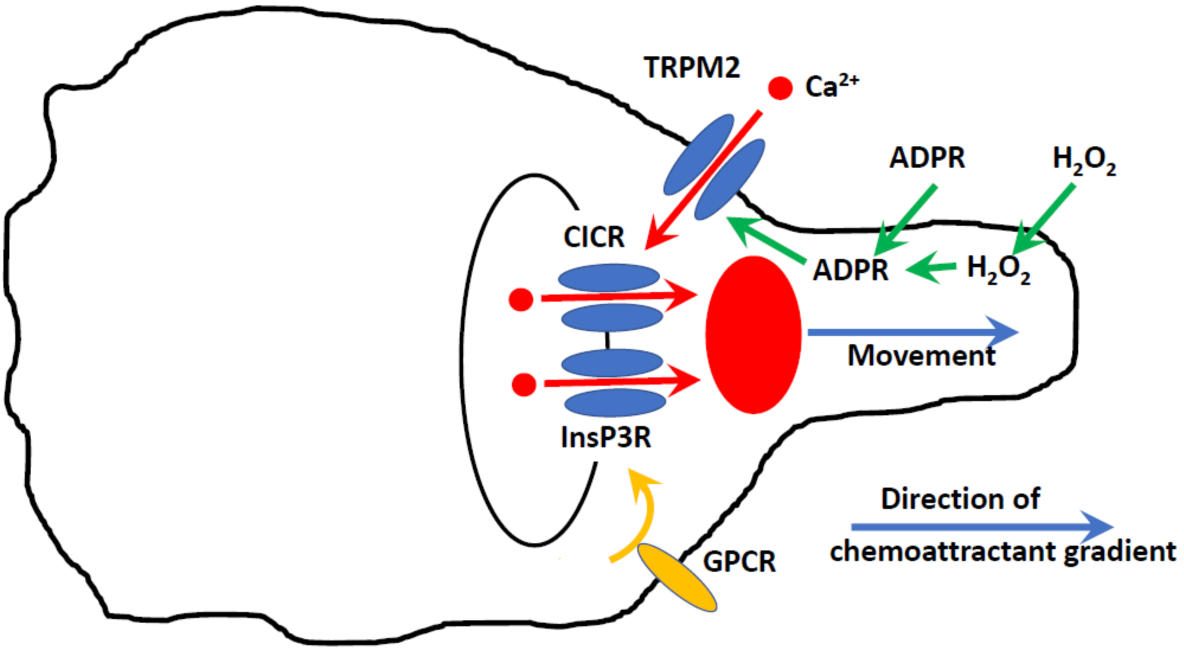


**Supplementary Materials Fig. 8. Summary diagram of proposed mechanism controlling neutrophil motility.**

The work described here supports the mechanism for activating neutrophil motility shown above. A gradient of $H_2O_2$ releases intracellular ADPR, preferentially activating calcium-permeable TRPM2 ion channels at the neutrophil leading edge. ADPR is also able to enter the cell directly, via an unknown mechanism. The calcium influx via TRPM2 is amplified by calcium-induced calcium release (CICR) from subcellular organelles. The resulting calcium "pulse" at the leading edge (red) activates the motile machinery, causing extension of pseudopodia in the direction of the $H_2O_2$ gradient.

Conventional chemoattractants such as LPS, C5a and CXCL2 bind to G-protein coupled receptors and also activate leading-edge calcium pulses by discharging intracellular calcium stores. A possible mechanism involves generation of IP3, followed by calcium release via the InsP3 receptor (InsP3R), though we have no direct evidence for this mechanism.

The intracellular signalling molecules Rac2 and PI3K modulate the amplitude of calcium pulses in all mechanisms and in this way modulate the motile machinery that produces pseudopodial extension. The intracellular signalling molecule *lyn* also modulates calcium release, but this affects only the pathway activated by $H_2O_2$, and has no effect on the amplitudes of Ca pulses nor movement in response to LPS, C5a or CXCL2.

### Supplementary Materials Fig. 9

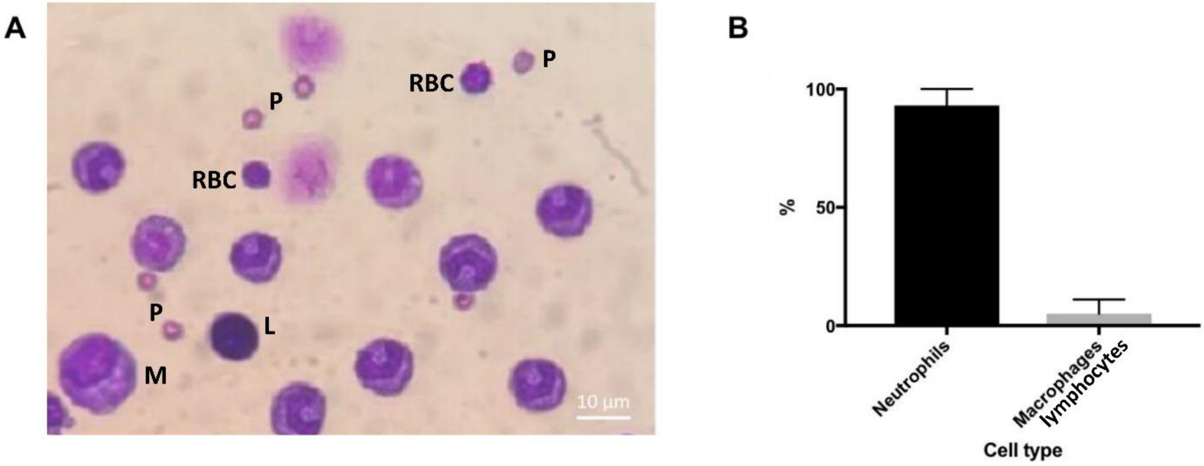


**Supplementary Materials Figure 9. Identification of cells from mouse peritoneal cavity.**

(A) Sample image containing major cell types observed in mouse peritoneal lavage following thioglycolate injection, prepared using Cytospin followed by a modified version of the May-Grünwald-Giemsa staining (see Methods). Neutrophils identified by their characteristic horseshoe-shaped nuclear morphology and size difference from macrophages. Lymphocytes identified by their large round nuclei. All cells visible in the image are neutrophils apart from red blood cells (RBC), platelets (P), one macrophage (M) and one lymphocyte (L). Note two degenerating neutrophils at top, identifiable from the horseshoe shapes of their nuclei. Scale bar bottom right.

(B) Neutrophils made up 94.25% ± 2.5% of the cells present in the peritoneal lavage, excluding non-motile cells. Macrophages and lymphocytes made up 5.75% ± 2.5% (n=4 individual experiments).
